# Efficient decoding of large-scale neural population responses with Gaussian-process multiclass regression

**DOI:** 10.1101/2021.08.26.457795

**Authors:** C. Daniel Greenidge, Benjamin Scholl, Jacob L. Yates, Jonathan W. Pillow

## Abstract

Neural decoding methods provide a powerful tool for quantifying the information content of neural population codes and the limits imposed by correlations in neural activity. However, standard decoding methods are prone to overfitting and scale poorly to high-dimensional settings. Here, we introduce a novel decoding method to overcome these limitations. Our approach, the Gaussian process multi-class decoder (GPMD), is well-suited to decoding a continuous low-dimensional variable from high-dimensional population activity, and provides a platform for assessing the importance of correlations in neural population codes. The GPMD is a multinomial logistic regression model with a Gaussian process prior over the decoding weights. The prior includes hyperparameters that govern the smoothness of each neuron’s decoding weights, allowing automatic pruning of uninformative neurons during inference. We provide a variational inference method for fitting the GPMD to data, which scales to hundreds or thousands of neurons and performs well even in datasets with more neurons than trials. We apply the GPMD to recordings from primary visual cortex in three different species: monkey, ferret, and mouse. Our decoder achieves state-of-the-art accuracy on all three datasets, and substantially outperforms independent Bayesian decoding, showing that knowledge of the correlation structure is essential for optimal decoding in all three species.

## 1 Introduction

Since Zohary, Shadlen, and Newsome’s landmark demonstration of correlated activity in a population of MT neurons (Zohary et al., 1994), computational neuroscience has been seeking to elucidate the role that noise correlations play in the population code (Adibi et al., 2013; Bartolo et al., 2020; Cafaro & Rieke, 2010; Ecker et al., 2011; Ecker et al., 2014; Kanitscheider et al., 2015; Kohn et al., 2016; Moreno-Bote et al., 2014; Nirenberg & Latham, 2003; Nogueira et al., 2020; Panzeri et al.,

Schneidman et al., 2003; Sokoloski et al., 2021). Noise correlations refer to statistical dependencies in the trial-to-trial fluctuations in population activity elicited in response to a fixed stimulus, and they may either increase or decrease information relative to a population with conditionally independent neurons (Averbeck et al., 2006; da Silveira & Rieke, 2020). A common strategy for evaluating the role that noise correlations play in a population code is to compare the accuracy of two decoders trained to predict stimuli from population activity: a “correlation-blind” decoder, which has no access to noise correlations present in population activity, and a “correlation-aware” decoder, which does (Berens et al., 2012; Bialek et al., 1991; Graf et al., 2011; Nirenberg & Latham, 2003; Pillow et al., 2008; Stringer et al., 2021; Yates et al., 2020). If the correlation-aware decoder performs better, then we may conclude that downstream regions must take noise correlations into account to optimally read out information from the upstream population.

This strategy is an effective way to investigate the scientific question, but existing work is plagued by a number of statistical issues, which we aim to address in this paper. First, as neural datasets have increased in dimensionality, regularization has become a prerequisite for good decoding performance, making it difficult to compare correlation-blind decoders—which are often fit without regularization—and correlation-aware decoders, which are almost always regularized. Second, conventional correlation-aware decoders struggle to scale statistically and computationally to modern datasets containing tens or hundreds of thousands of neurons.

To address these shortcomings, we develop a suite of three new decoders with a common regularization strategy based on Gaussian Processes (GPs). First, we introduce two correlation-blind decoders that apply Bayesian decoding to an independent encoding model: the GP Poisson Independent Decoder (GPPID), which assumes independent Poisson encoding noise; and the GP Gaussian Independent Decoder (GPGID), which assumes independent Gaussian encoding noise. Both of these decoders place a Gaussian process prior over the neural tuning curves. (For each neuron, its tuning curve is its mean response as a function of the stimulus variable.) The GPPID model can be used when the neural responses are encoded by non-negative integers (e.g., spike counts), whereas the GPGID model can be used when the neural responses are real numbers (e.g., calcium imaging). We emphasize that both of these decoders are insensitive to correlations in neural activity, because they rely on independence assumptions.

We then introduce a novel correlation-aware decoding model, the Gaussian Process Multiclass Decoder (GPMD), which is a multinomial logistic regression model that uses a GP prior to regularize the decoding weights for each neuron. This decoder, which learns a direct linear mapping from high-dimensional neural activity patterns to the log-probability of the stimulus, is the only one of the three that take noise correlations into account. However, the three decoders have a similar number of parameters—equal to the number of neurons times the number of stimulus categories—and rely on a common regularization method, making it straightforward to compare them.

We compared our decoders to a variety of previously proposed decoding methods: first, multinomial logistic regression regularized using an elastic-net penalty (GLM-NET, see Zou and Hastie [2005]); second, the empirical linear decoder (ELD), a decoder trained using support vector machines (Graf et al., 2011); and third, the “super-neuron” decoder (SND), a recently proposed decoder trained using least squares regression and a bank of nonlinear target functions (Stringer et al., 2021). All three of these decoders are correlation-aware linear classifiers. For completeness, we also compared our decoders to unregularized, correlation-blind Poisson and Gaussian independent decoders (PID/GID).

We benchmarked all these decoders on three real-world datasets from primary visual cortex (V1), recorded from monkey (Graf et al., 2011), ferret (this paper), and mouse (Stringer et al., 2021). We found that our regularized correlation-blind decoders (GPPID and GPGID) could match and even exceed the performance of some of the correlation-aware decoders. However, none of these decoders performed as well as our our proposed correlation-aware decoder, the GPMD, which achieved state-of-the-art accuracy on all datasets, with no preprocessing or manual elimination of noise neurons. The GPMD’s performance advantage derived directly from its expressive regularization strategy, which successfully adapted to every dataset’s unique structure while remaining as scalable as other regularized, correlation-aware decoders.

These results indicate that knowledge of the correlation structure is crucial for optimal readout of stimulus information from V1 populations in all three species. For ease of use, our decoders conform to the scikit-learn interface and are released as a Python package at https://github.com/cdgreenidge/gdec.

## 2 The neural decoding problem

In this paper, we consider the problem of a decoding a low-dimensional stimulus variable (i.e., the orientation of a sinusoidal grating) from a high-dimensional neural activity pattern (i.e., a vector of spike counts). We assume the stimulus belongs to one of *K* discrete bins or classes, formally making this a classification problem. However, the regression problem can be approximated by making *K* large, so that the grid of stimulus values becomes arbitrarily fine.

Figure 1 illustrates the problem setup for the V1 datasets we examined. The visual stimulus for each individual trial is a drifting sinusoidal grating with an orientation *θ_k_* selected from a set of discrete orientations {*θ*_1_, …, *θ_K_*} that evenly divide the interval [0, 2*π*]. The stimulus variable to be decoded is thus a categorical variable *y* ∈{1, …, *K*}.

**Figure 1:**
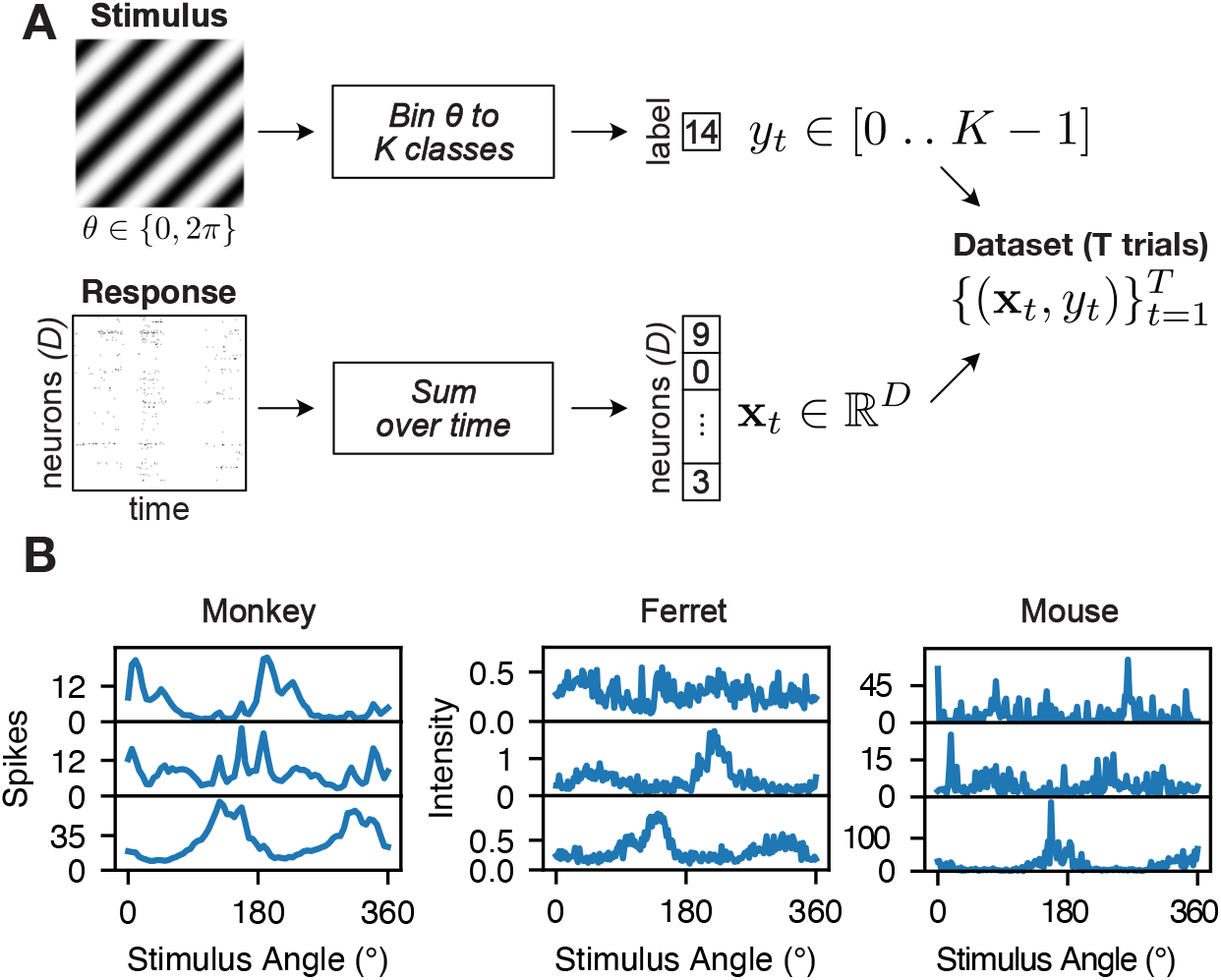
**A**: Decoding task diagram. The animal is presented a grating drifting at an angle *θ* ∈ [0, 2*π*]. Responses are recorded from primary visual cortex and summed over time into a feature vector, **x**. The stimulus is binned to an integer class label, *y*. We use linear decoders to predict *y* from **x**. **B**: Tuning curves from three randomly selected neurons from each species. The two calcium fluorescence datasets (ferret and mouse) are noisier and have many more neurons, increasing the importance of regularization.

We consider the neural population response to be a vector 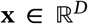, where *D* indicates the number of neurons in the dataset. We obtained this response vector by summing each neuron’s spikes (monkey) or two-photon calcium fluorescence (ferret and mouse) over some time window after stimulus presentation. Figure 1B shows orientation tuning curves from three example neurons from each dataset. The monkey datasets (left) contained between *D* = 68 and *D* = 147 neurons, with *K* = 72 discrete stimulus orientations (spaced every 5 degrees), and 50 trials per orientation for *T* = 3600 trials (Graf et al., 2011). The ferret dataset (middle) contained *D* = 784 neurons, with *K* = 180 discrete stimuli (spaced every 2 degrees) and 11 trials per orientation for a total of *T* = 1991 trials, with the 0°/360° orientation sampled twice. Finally, the mouse datasets (right) contained between *D* = 11311 and *D* = 20616 neurons. The stimuli for this experiment were sampled uniformly in [0, 2*π*], and we subsequently discretized them into *K* = 180 bins (Stringer et al., 2021). Each bin contained between 12 and 42 trials for a total of between *T* = 4282 and *T* = 4469 trials, depending on the dataset.

In each case, we collected the population response vectors **x** and the discretized stimuli *y* into a classification dataset 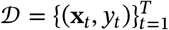. Full details on these datasets and their preprocessing procedures can be found in Appendix C.

### 2.1 Linear classification

The decoders we consider are all linear classifiers, meaning that they are defined by a set of linear decoding weights and an intercept term. Their common form is:

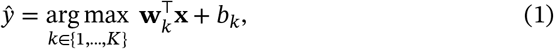

where 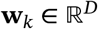 is a set of decoding weights, and *b_k_* is an intercept term for stimulus class *k*. Note that an explicit intercept term is not strictly necessary, since it can be included in the weights if a 1-valued entry is appended to **x**. To obtain the stimulus estimate 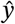, we compute the dot product between the neural response vector **x** and the weights for each class, and select the class in {1, …, *K*} for which this dot product is maximal.

The full set of parameters for a decoding model is thus the set of decoding weights for each class, which can be written as a *D* × *K* matrix *W* = (**w**_1_, …, **w**_K_), and, optionally, a *k*-dimensional intercept vector **b** = (*b*_1_, …, *b_k_*)^⊤^. In the following, we will let *W_dk_* denote the decoding weight for neuron *d* for stimulus class *k*. The only difference between the decoding methods we will consider is the procedure for training these weights from data.

The probabilistic formulation of these classifiers is that the log-probability of the stimulus class is an affine (i.e., linear plus constant) function of the neural response:

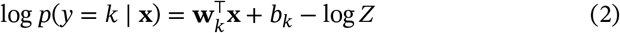

or equivalently

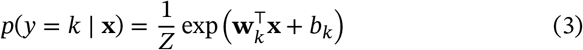

where 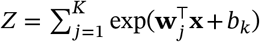 is the normalizing constant ensuring that the probabilities over the *K* classes sum to 1.

Decoding with linear classifiers is optimal for so-called exponential-family probabilistic population codes with linear sufficient statistics (Beck et al., 2007; Ma et al., 2006). Although we could certainly expand our study to consider decoding with nonlinear classifiers, previous analyses of two of the datasets we used showed no benefit from adding nonlinear classification (Graf et al., 2011; Stringer et al., 2021).

### 2.2 Noise and signal correlations

It is helpful to distinguish between two different types of correlations present in neural population responses: noise correlations and signal correlations (Averbeck et al., 2006; Panzeri et al.,). Noise correlations correspond to statistical dependencies in *P*(**x** | *y*), the distribution of population responses **x** given a stimulus *y*. If neurons are conditionally independent (as assumed by the Poisson and independent Gaussian encoding models), then noise correlations are zero. Noise correlations are typically characterized by Cov[**x** | *y*], the covariance of the population responses to a fixed stimulus. This covariance clearly depends on the stimulus, meaning that noise-correlations may differ for different stimuli.

Signal correlations, on the other hand, are correlations between the tuning curves of different neurons. Two neurons with similar tuning curves will have high signal correlations, while neurons with dissimilar tuning curves may have zero or even negative signal correlations (i.e., if their tuning curves are anti-correlated). Signal correlations are commonly characterized by the covariance matrix 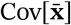, which is the covariance of the tuning curves 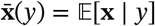 over the stimulus distribution *P*(*y*).

The effects of noise correlations on decoding performance depend critically on the alignment between noise and stimulus correlations Averbeck and Lee (2006), da Silveira and Rieke (2020), and Panzeri et al. (). However, it is important to note that so-called “correlation-blind” decoders are blind only to noise correlations. Thus, comparing correlation-blind and correlation-aware decoders is a method for determining whether a downstream population needs to know the structure of the noise correlations in order to decode information optimally. Figure S1 shows a characterization of both noise and signal correlations for the datasets considered in this paper.

## 3 Review of existing decoders

Here we describe previously proposed neural decoding methods, to which we will compare the Gaussian Process-based decoders which will be introduced in Section 4.

### 3.1 Correlation-blind decoders

First, we introduce two independent or “correlation-blind” decoders, the first assuming Poisson noise, and the second assuming Gaussian noise. Both decoders use Bayes’ rule to obtain a posterior distribution over the stimulus under an independent encoding model, an approach commonly known as “naïve Bayes.” The encoding models underlying these decoders assume that neural responses are conditionally independent given the stimulus, preventing them by construction from using noise correlations for decoding.

#### 3.1.1 The Poisson Independent Decoder (PID)

The Poisson independent decoder relies on an independent Poisson encoding model of neural responses, which assumes that each neuron’s spike count is drawn from an independent Poisson distribution the mean of which is determined by the stimulus (Abbott, 1994; Földiák, 1993). The encoding model describes *x_d_*, the response of neuron *d*, as:

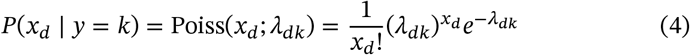

where *λ_dk_* is the mean response of neuron *d* to stimulus *k*. Under the conditional independence assumption, the joint distribution of the population response is the product of the single-neuron encoding distributions:

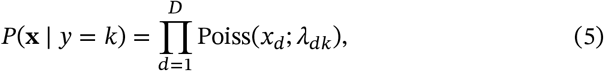

where *D* is the total number of neurons.

Using Bayes’ theorem, it is possible to derive the probability over stimuli given an observed response vector **x**:

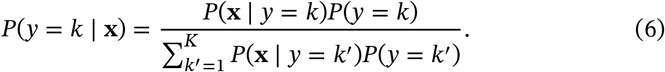

This quantity can be used for decoding. If there are equal numbers of trials per class in the training dataset, the prior probabilities *P*(*y* = *k*) are equal for all *k*, and cancel, leaving the prediction rule

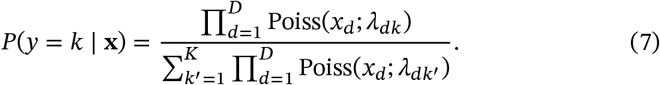

To fit the model, one must estimate the parameters {*λ_dk_*} for every stimulus and neuron. The vector of parameters for a single neuron ***λ**_d_* = (*λ*_*d*1_, …, *λ_dK_*)^T^ is known as the tuning curve. It contains the neuron’s expected response as a function of the stimulus. The maximum likelihood estimate for *λ_dk_* is given by the mean spike count for each neuron-stimulus combination:

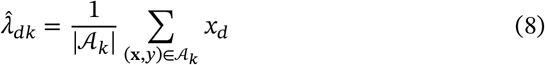

where 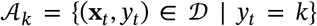 is the set of all elements of the dataset 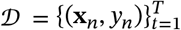 associated with a particular stimulus *y* = *k*, and 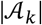 is the number of elements in 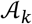.

Assuming that the prior class probabilities *P*(*y* = *k*) are equal, the log of the class-conditional probability (eq. 7), also known as the log posterior over classes, can be written:

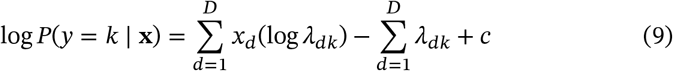

where *c* is a constant we ignore because it does not depend on the class. This shows that the PID decoder is a linear classifier (eq. 1), with weights 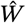 and intercepts 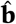 given by

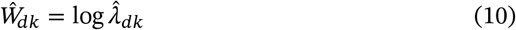

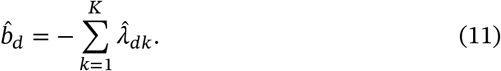

See Appendix A for a detailed derivation.

#### 3.1.2 The Gaussian Independent Decoder (GID)

The Poisson independent decoder described above can only applied to nonnegative integer data, such as spike counts. For real-valued data such as calcium fluorescence, intracellularly-recorded membrane potential, local field potential, or fMRI BOLD signals, it is common to use a Gaussian encoding model. This model describes *x_d_*, the response of neuron *d*, as:

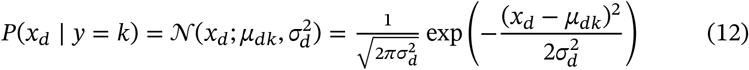

where *μ_dk_* is the mean response of neuron *d* to stimulus *k*, and 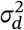 is the noise variance for neuron *d*. Unlike a typical Gaussian naïve Bayes decoder, we restrict the noise variance to be constant across stimulus classes, though, as usual, it can vary across neurons. With this restriction, the decoder becomes a linear classifier, like the other decoders we consider. If the noise variance were allowed to vary across stimulus classes, the decoder would be a quadratic classifier (see Appendix A.)

To fit the model, we compute maximum likelihood estimates of the encoding distribution parameters for each neuron, which are given by the class-conditional empirical means 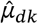, and the empirical variances 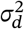, for each *d*-th neuron:

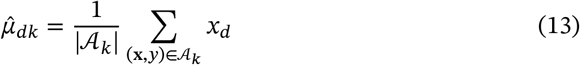

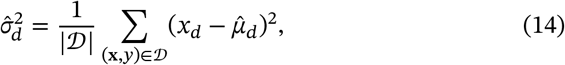

As before, 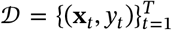 is the dataset, and 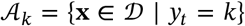 is the set of all neural response vectors for a particular stimulus *y* = *k*.

Decoding stimuli under this encoding model follows from Bayes’ rule in the same manner as in the Poisson independent decoder (eq. 7), but using the Gaussian encoding distribution instead of the Poisson. After some algebra, we can see that the Gaussian independent decoder (GID) is a a linear classifier (eq. 1) with weights *W* and intercepts **b** given by

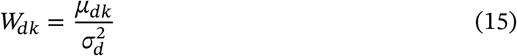

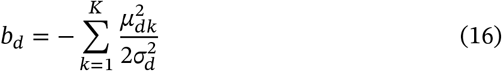

See Appendix A for a detailed derivation.

### 3.2 Correlation-aware decoders

Here we review three previously-proposed decoders that take into account the structure of neural correlations when determining a classification boundary, and are therefore “correlation-aware.” Unlike the two naïve Bayes decoders described above, which resulted from applying Bayes’ rule to an encoding model, these directly model the posterior probability over stimuli given a vector of neural activity. All three decoders are multiclass linear classifiers, but they are trained with different loss functions and regularization methods.

#### 3.2.1 Multinomial logistic regression with elastic-net penalty (GLMNET)

The multinomial logistic regression model is a generalization of binary logistic regression to the multiple-class setting. It assumes that the log probability of the stimulus given the response is an affine function—that is, a linear transform plus a constant—of the neural response vector (Bishop, 2006). The conditional probability of the stimulus belonging to class *k* given the neural response vector **x** can be written:

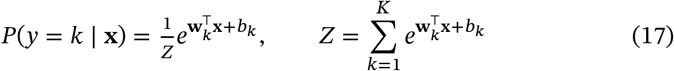

where **w**_*k*_ is a vector of the decoding weights for class *k*, *b_k_* is the constant offset for class *k*, and the *Z* is the normalizing constant.

The model parameters consist of the weights *W* = (**w**_1_, …, **w**_*K*_) and offsets **b** = (*b*_1_, …, *b_K_*)^⊤^, and can be fit by maximum likelihood. The log-likelihood function given the dataset 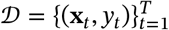 can be written:

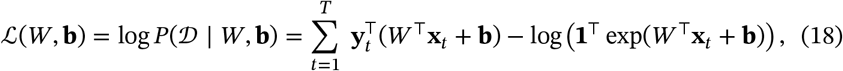

where **y**_*t*_ is the one-hot vector representation of the stimulus class *y_t_* ∈ {1, …, *K*} on trial *t*—that is, a vector of all zeros except for a one in the entry corresponding to the stimulus class—and **1** is a length-*D*) vector of ones.

The maximum-likelihood estimator (MLE) tends to perform poorly in settings with limited amounts of data, and may not exist for small datasets. In fact, the MLE is not defined when the number of trials *T* is smaller than the number of identifiable parameters in the weight matrix *W*, i.e. when *T* < *D*)(*K* – 1). Even in settings where the MLE does exist, it may overfit, yielding poor generalization performance.

A popular solution to this problem is to regularize the MLE with the elastic-net penalty, which combines ℓ_1_ (“lasso”) and ℓ_2_ (“ridge”) penalties to induce parameter sparsity and shrinkage (Friedman et al., 2010). The elastic-net estimator is obtained by maximizing the log-likelihood minus the regularization penalty:

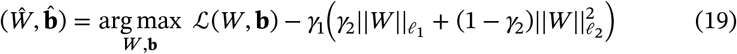

Here, *γ*_1_ is a hyperparameter determining the strength of the regularization, and *γ*_2_ is a hyperparameter that controls the balance between the ℓ_1_ penalty, which encourages *W* to be sparse, and the ℓ_2_ penalty, which encourages *W* to have a small magnitude. For our decoding tasks, we found that including the ℓ_2_ penalty always diminished cross-validated performance, so we fixed *γ*_2_ = 0. We then set *γ*_1_ using a five-step logarithmic grid search from 10^−4^ to 10, evaluated with three-fold cross validation.

#### 3.2.2 The Empirical Linear Decoder (ELD)

The Empirical Linear Decoder (ELD), introduced by Graf et al. (2011), is similar to multinomial logistic regression in that it models the log probability of the stimulus class as an affine function of the neural response vector (eq. 17). However, instead of using standard likelihood-based methods to fit the model, the authors constructed an inference method based on support vector machines (SVMs).

Their key observation was that the log-likelihood ratio for adjacent stimulus classes is an affine function of the response vector, with weights given by the difference of the two classes’ decoding weight vectors. For example, for stimulus classes one and two we have:

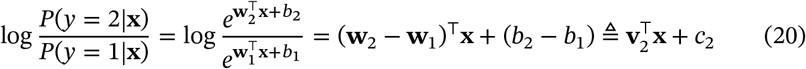

Here, we have defined **v**_2_ to be the difference vector (**w**_2_ – **w**_1_) and *c*_2_ to be the difference scalar *b*_2_ – *b*_1_.

We see that discriminating class two from class one under the multinomial logistic regression model is equivalent to solving a linear binary classification task with weights **v**_2_ and offset *c*_2_. The authors proposed estimating **v**_2_ and *c*_2_ using an SVM trained on the data from classes one and two. They then used the same approach to estimate the weights for all subsequent pairs of adjacent classes. That is, they estimated the difference weights **v**_*k*+1_ using an SVM trained on data from classes *k* – 1 and *k*, for *k* = 2, …, *K*.

To recover the weights of the multinomial logistic regression model from the SVM weights, Graf et al. (2011) used the recursions:

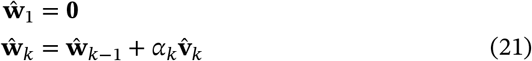

and

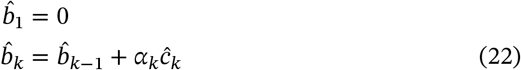

for *k* = 2, …, *K*. Here the weights for class one can be set to zero without loss of generality. The constant *α_k_*, which scales the contribution of the SVM weights **v**_k_ and *c*_k_, is necessary because SVMs only recover **v**_*k*_ and *c_k_* up to a multiplicative constant. We were unable to determine how the authors set these scaling constants (Figure S6), so we fit them by maximizing the log likelihood of the data under the multinomial logistic regression model (eq. 18).

#### 3.2.3 The Super Neuron Decoder (SND)

The Super Neuron Decoder (SND), introduced by Stringer et al. (2021), is a third approach for training a linear classifier on multi-class data. It optimizes a set of decoding weights using penalized least-squares regression and a set of nonlinear super-neuron response functions. The super-neuron response functions encode the tuning curves of a population of narrowly-selective downstream “super-neurons”, containing one super-neuron for each stimulus orientation. Each super-neuron responds maximally to a single orientation, making the population response on each trial a narrow bump of activity centered on the correct stimulus.

Formally, the SND seeks a matrix of weights *W* that maps the response vector **x** to a target vector 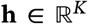 on each trial. The target vector **h** contains the responses of the super-neuron population. The super-neurons have tuning curves parameterized in the same manner as the von Mises probability density function, which is appropriate since the stimulus variable is periodic.

The *i*-th super-neuron has a preferred orientation of 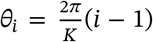, so its tuning curve is given by:

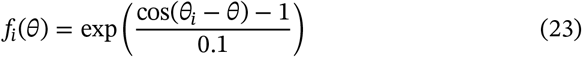

The target vector for the *k*-th stimulus class is therefore

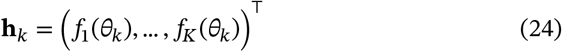

where 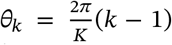 is the stimulus angle associated with stimulus class *k* ∈ {1, …, *K*}.

Stringer et al. (2021) trained the model weights *W* by linearly regressing the observed neural responses onto the target vectors. To penalize large weight values, they included an ℓ_2_ (“ridge”) regularization:

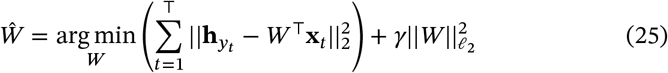

The term 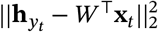 is the squared error between the correct target vector **h**_*y_t_*_ and the output of the linear decoding weights *W*^⊤^**x**_*t*_ on trial *t*. The term 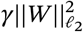 is the squared ℓ_2_ penalty on the decoding weights with regularization strength *γ*, which the authors fixed at *γ* = 1.0. Intuitively, this training procedure seeks weights *W* that make the linearly transformed population response *W*^⊤^**x** match the superneuron population response **h** as closely as possible in a least-squares sense.

The decoding rule chooses the class label corresponding to the maximum of the linearly weighted responses. (This is the same decoding rule as in the other decoders we have considered):

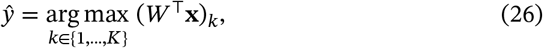

Here (*W*^⊤^**x**)_*k*_ is the *k*-th element of the transformed response vector *W*^⊤^**x**. In other words, the predicted stimuli value is the preferred orientation of the maximally responding super-neuron.

## 4 Proposed methods: GP-regularized decoders

In this section, we first introduce two correlation-blind decoders regularized with Gaussian process (GP) priors: the GP-regularized Poisson Independent Decoder (GPPID) and the GP-regularized Gaussian Independent Decoder (GPGID). After that, we will introduce a correlation-aware decoder, the Gaussian Process Multiclass Decoder (GPMD), which adds a GP prior to multinomial logistic regression for the same purpose.

Gaussian Processes are often used for nonlinear classification problems, a method setting known as Gaussian Process classification (Bishop, 2006; Liu et al., 2021). In neuroscience settings, GPs have also been used to construct priors over smoothly-varying firing rates or latent factors (Cunningham et al., 2008; Duncker & Sahani, 2018; Keeley et al., 2020; Yu et al., 2009). Here, we use GPs to provide a construct over the weights in a general linear model (in the case of GPGID) or generalized linear model (GPPID and GPMD). Previous work has used GPs to provide a smoothing prior overweights of a linear or generalized linear model (Macke et al., 2011; Park & Pillow, 2011; Park et al., 2014; Rad & Paninski, 2010; Sahani & Linden, 2002). However, to our knowledge, ours is the first to use GP priors to impose a smoothness assumption over the different classes in a multinomial logistic regression model.

In the following sections, we will describe the relationship between the linear weights in our model and neural tuning curves, which makes GPs a natural tool for regularization in our problem.

### 4.1 The GP-regularized Poisson Independent Decoder (GPPID)

When doing inference in the PID decoder (section 3.1.1), it is necessary to estimate *λ_dk_*, the expected spike count of neuron *d* in response to the *k*-th stimulus, across all neurons and stimuli. The expected spike counts for a single neuron forms a tuning curve across stimuli, ***λ**_d_* = (*λ*_*d*1_, …, *λ_dK_*)^*T*^. The maximum likelihood estimator for each entry in the tuning curve is simply the empirical mean of the spike counts for the *d*-th neuron under the *k*-th stimulus. However, the empirical mean estimates are noisy, especially when the number of trials for each stimulus is small, which can limit the PID decoder’s performance.

In principle, we could compensate for the noise by recording more trials for each stimulus, but this is expensive, particularly if the stimulus grid has a fine resolution. Instead, we propose to reduce error in the tuning curve estimates by exploiting our prior knowledge that tuning curves tend to be smooth with respect to orientation. We incorporate this knowledge into the model by placing an independent Gaussian process prior over the log tuning curve of each neuron (Park et al., 2014; Rad & Paninski, 2010).

The resulting GP-regularized PID model is given by

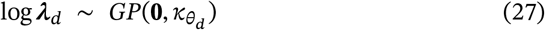

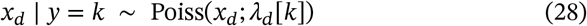

where *x_d_* is the spike response of neuron *d*, and *λ_d_*[*k*] is the *k*th element of the tuning curve ***λ***_*d*_. Here, the log tuning curve has a Gaussian Process prior with zero mean, and a covariance function *κ*(·, ·) with hyperparameters *θ_d_*.

#### 4.1.1 Periodic Gaussian covariance function

Because the stimulus orientations live on a circular domain, we selected a periodic covariance function for the GP prior. In particular, we used covariance based on the Gaussian covariance function, also known as the radial basis function (RBF) or squared-expoential covariance function. To make this covariance periodic, this covariance is wrapped around the circle as follows (Scholkopf & Smola, 2002):

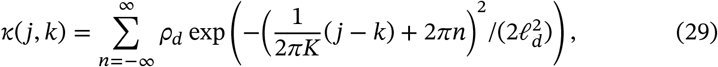

where indices *j*, *k* ∈ {1, … *K*} denote stimulus classes, and the hyperparameters *θ_d_* = {*ρ_d_*, ℓ*_d_*}, are the marginal variance *ρ_d_* and length scale ℓ*_d_* of the log tuning curve. The sum over integer multiples of 2*π* accomplishes wrapping, making the function periodic. When the length scale ℓ*_d_* is much smaller than 2*π* this function can often be evaluated accurately using a single term from the infinite sum. However, our implementation relies on a sparse Fourier-domain implementation, meaning that we do not need to evaluate it in the orientation domain at all. (See Appendix B for details).

The neuron-specific hyperparameters *θ_d_* permit tuning curves to differ in amplitude and smoothness, so different neurons can be regularized differently. For neurons with non-existent tuning or exceptionally noisy responses, the inference procedure will set the amplitude *ρ_d_* to zero or the length scale ℓ*_d_* to infinity, making the tuning curve flat (see Figure 5). Such neurons are effectively pruned from the dataset, since flat tuning curves make no contribution to decoding. This effect is known as automatic relevance determination (MacKay, 1992; Neal, 1996), and it eliminates the need to manually filter out noisy or untuned neurons, which can be critical when using other decoding methods (Graf et al., 2011).

#### 4.1.2 Fitting the GPPID to spike count data

To fit the GPPID model to spike count data, we employ a two-step procedure known as empirical Bayes (Bishop, 2006). For each neuron, we first compute a point estimate of the neuron-specific tuning curve hyperparameters by maximizing the model evidence, also known as marginal likelihood. Then, we find the *maximum-a-posteriori* (MAP) estimate of the tuning curve using the estimated hyperparameters 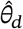. The model evidence for neuron *d* is the marginal probability of **X**_\**d*_ = (*x*_1*d*_, …, *x_Td_*)^⊤^, the *d*-th column of the spike count matrix **X**, given the hyperparameters *θ_d_* and stimuli *Y*:

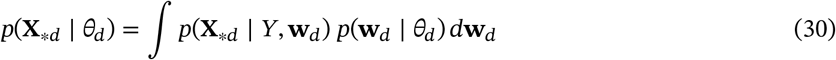

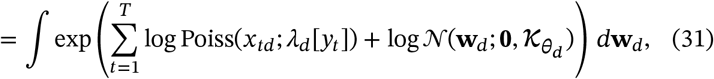

where **w**_*d*_ = log ***λ**_d_* is the log-tuning curve for neuron *d*, ***λ***_*d*_[*y_t_*] denotes the element of ***λ**_d_* = exp(**w**_*d*_) corresponding to *y_t_*, the stimulus class for the *t*-th time bin, and *κ_θ_d__* denotes the *K* × *K* covariance matrix arising from evaluating the periodic Gaussian covariance function (eq. 29) at all pairs of points in (1, …, *K*).

Because the integral in (eq. 31) is intractable, we use the standard approach of approximating it using Laplace’s method (Bishop, 2006). In the following, let 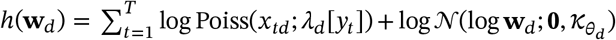 denote the sum of log-likelihood and log-prior, and *H* be the Hessian (second-derivative matrix) of *h*(**w**_*d*_), evaluated at its maximizer, the MAP estimate 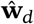. The approximate evidence is then given by:

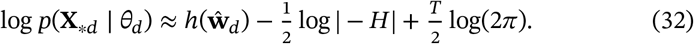

We compute point estimates of the model hyperparameters 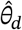 by optimizing this approximation with respect to *θ_d_* using the Nelder-Mead algorithm (Nelder & Mead, 1965).

Once we have estimated the maximum-evidence estimate of the hyperparameters 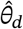 for neuron *d*, we compute the empirical Bayes estimate of the neuron’s logtuning-curve by computing the MAP estimate of **w**_*d*_ under the model given by (eqns. 27-28):

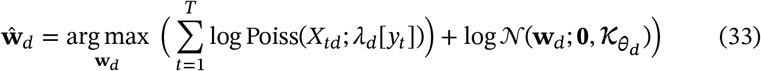

where Poiss(*X_td_*;***λ**_d_*[*y_t_*]) is the probability of spike count *X_td_* given the firing rate ***λ**_d_*[*y_t_*] = exp(**w**_*d*_[*y*_*t*_) under a Poisson distribution (eq. 4).

To accelerate the optimization procedure, we use a Fourier-domain representation of the covariance matrix *κ_θ_d__* based on the Karhunen-Loéve expansion (Paciorek, 2007; Royle and Wikle, 2005; Wikle, 2002; see Appendix B for details.). Since the procedure described above can be performed independently for each neuron, fitting the GPPID model is fully parallelizeable across neurons.

### 4.2 The GP-regularized Gaussian Independent Decoder (GPGID)

In the same way that we added a Gaussian process prior to the Poisson independent decoder to obtain the GPPID decoder, we can add a GP prior to the Gaussian independent decoder to obtain a GP-regularized Gaussian independent decoder (GPPID). However, three salient differences set the Gaussian model apart from the Poisson case. First, the parameters of interest in the Gaussian model are the tuning curves, not the log-tuning curves that appeared in the Poisson case above. Second, there is an additional hyperparameter to learn in the Gaussian case, corresponding to each neuron’s observation noise 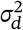. Third, and most importantly, the Gaussian model evidence can be computed in closed form, obviating the need for approximations.

The GPGID model for the *d*’th neuron is defined by a GP prior over the tuning curve ***μ**_d_* = (*μ*_1*d*_, …, *μ_Kd_*)^*T*^ and a Gaussian observation model:

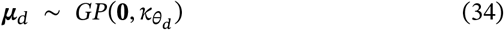

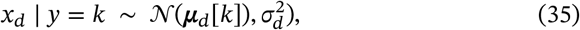

where *x_d_* is the response of neuron *d, **μ**_d_* [*k*] is the *k*th element of the tuning curve ***μ**_d_*, and *κ_θ_d__* is the periodic Gaussian covariance function defined in (eq. 29). The model contains three hyperparameters for each neuron, 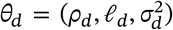, consisting of the marginal variance *ρ_d_* and length scale ℓ_*d*_ of the GP prior, and the observation noise variance 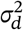, which appears in the likelihood. As in the GID model (section 3.1.2), we ensure that the naïve Bayes classifier is linear by assuming that the observation noise variance is constant over classes (see Appendix A).

To fit the model, we again use empirical Bayes. The first step is thus to compute a maximum-evidence estimate of the hyperparameters *θ_d_* for each neuron:

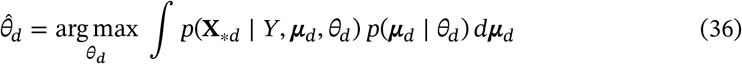

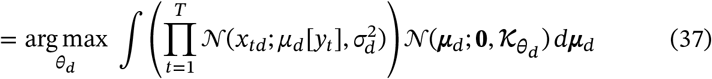

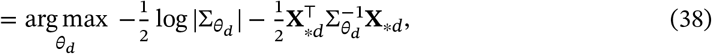

where *μ_d_*[*y_t_*] represents the tuning curve value for *y_t_*, the stimulus class on trial *t*, *κ_θ_d__* is the *K* × *K* covariance matrix induced by the GP prior, and 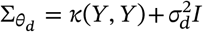 denotes a *T* × *T* covariance matrix whose *t*’th element along the main diagonal is 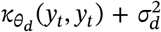 and off-diagonal element (*s,t*) is equal to *κ_θ_d__*(*y_s_, y_t_*). Because this objective function can be expressed analytically, we perform the maximization using a trust-region Newton method.

Once we have optimized (eq. 38) to find 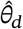, we compute the MAP estimate of the tuning curve given these hyperparameters, which has an analytic solution given by:

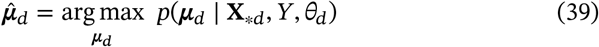

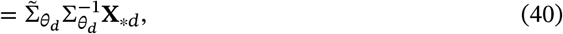

where 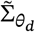 is a *K*×*T* matrix with *k*, *t*’th element given by *κ_θ_d__*(*k*, *y_t_*), for *k* ∈ (1, … *K*) (Rasmussen & Williams, 2006). However, for scalability we use an equivalent formula leveraging the same spectral weight representation as in the GPPID. (See Appendix B for details.)

### 4.3 The Gaussian Process Multiclass Decoder (GPMD)

In this section, we introduce the the Gaussian Process Multiclass Decoder (GPMD), which is multinomial logistic regression with a GP prior placed over the weights for each neuron (see Figure 2). As in section 3.2.1, the multinomial logistic regression model can be written:

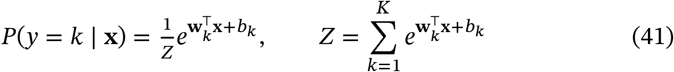

where **w**_*k*_ is a vector containing the decoding weights for class *k*, *b_k_* is the offset for class *k*, and *Z* is the normalizing constant.

**Figure 2:**
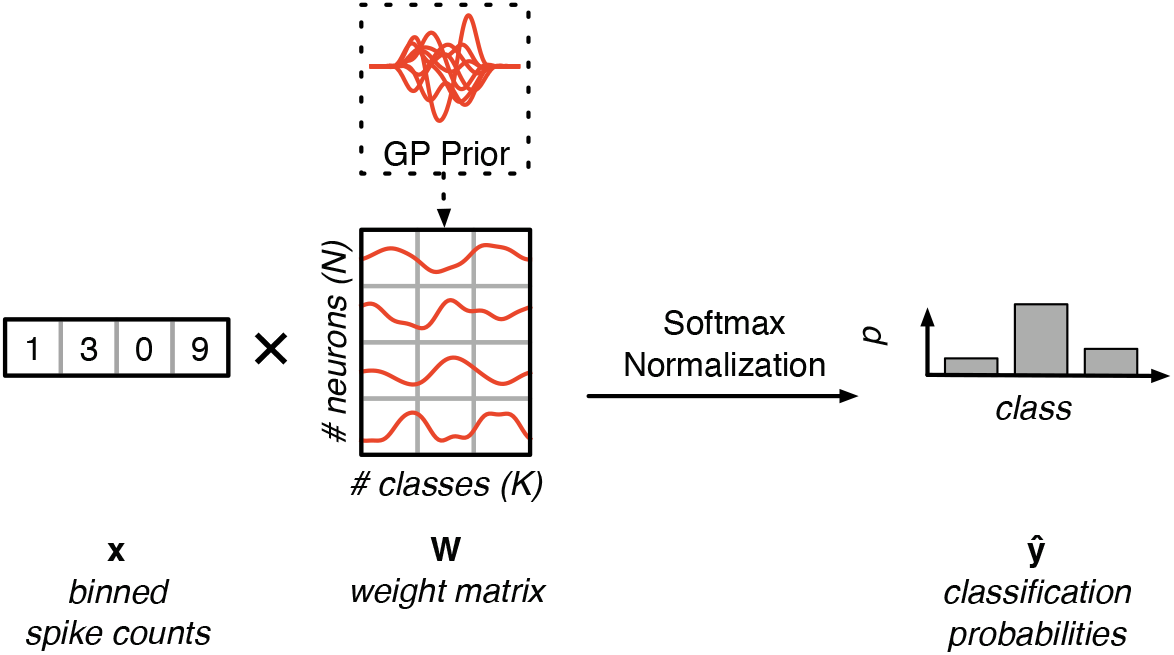
The GPMD model. A smoothing Gaussian Process prior is applied to each row of a logistic regression weight matrix **W** to shrink the weights and to ensure that they vary smoothly across stimuli, mimicking the smoothness of neural tuning curves. The weight matrix is then linearly combined with the neuron’s response vector **x** to produce unnormalized scores for each decoding class. A softmax function is used to transform the unnormalized scores into probabilities.

We regularize the weight matrix by placing an independent zero-mean Gaussian Process prior on each of its rows:

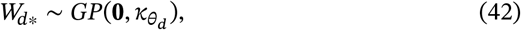

Here *W_d*_* is the *d*-th row of *W*, which contains the decoding weights associated with neuron *d* across all stimuli, and *κ_θ_d__* is the periodic Gaussian covariance function defined in (eq. 29).

In the GPPID and GPGID, we used a GP prior to formalize our prior knowledge that neuron’s log-tuning curves, and thus its tuning curves, tend to be smooth. In the GPMD, the model weights have no direct interpretation in terms of tuning curves. However, we can still motivate the use of a GP prior by observing that decoding weights also ought to vary smoothly with orientation, since tuning varies smoothly as a function of orientation.

Like previous decoders, the GPMD has neuron-specific hyperparameters, which allow different neurons to have decoding weights with different amplitudes and different amounts of smoothness. This flexibility has two benefits: first, it allows each neuron’s weights to adapt to the neuron’s response properties, and second, it automatically discards untuned or noisy neurons as described in section 4.1, eliminating the need for manual dataset preprocessing.

To fit the GPMD, we use variational inference to simultaneously learn both a posterior estimate for the weights {*W*, **b**} and point estimates for the prior hyperparameters. Specifically, given an approximate posterior family *q*—which we choose to be mean-field Gaussian—indexed by parameters *ϕ* and prior hyperparameters *θ*, we maximize the evidence lower bound (ELBO, see Blei et al. [2017] and Hoffman et al. [2013]) jointly with respect to *ϕ* and *θ*:

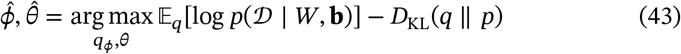

To calculate the likelihood term, we draw *M* = 3 samples from the variational posterior 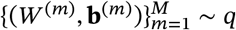, and use these to compute a Monte Carlo approximation of the expectation:

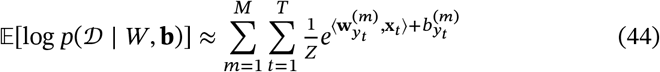

where *Z* is the normalizing constant defined in Eq. 41. In principle, we could also calculate this approximation using a subset of the data, or “minibatch” (Hoffman et al., 2013), but our datasets are small enough that this is not necessary, and doing so increases the approximation variance.

The KL-divergence term expands to a sum of KL divergences for each row of the weight matrix, since each row of the matrix is independent of the others. Assuming *q*_1_, …, *q_D_* are the mean-field variational distributions for each row of the weight matrix, and *p*_1_, …, *p_D_* are the associated GP priors, the KL-divergence term reduces to

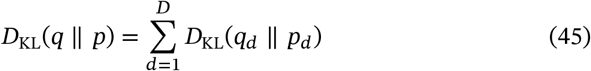

Each of the summands is easy to calculate analytically, since *q_d_* and *p_d_* are both multivariate normal distributions. The approximate variational posterior *q_d_* is a diagonal normal distribution, and the GP prior for each column *p_d_* is the multivariate normal distribution 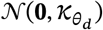, where, as in the GPPID model, *κ_θ_d__* is the *K* × *K* prior covariance matrix induced by the GP prior.

To make predictions, we approximate MAP inference by using the mode of the posterior approximation *q* as a point estimate of the weights *W* and **b** in the multinomial logistic regression model (eq. 41). Since we are only interested in the effects of the prior, and not in using the posterior to assess prediction uncertainty, this approach suffices. In practice, we found that setting of the offset **b** to 0 made no significant difference in performance (Figure S7), likely because all our datasets were reasonably high-dimensional.

#### 4.3.1 Scaling GPMD inference

When maximizing the ELBO, there are two scaling concerns: the number of examples (*T*) and the number of neurons (*D*). It is well-known that the ELBO can be scaled to huge numbers of examples by estimating the likelihood term using a minibatch approximation (Hoffman et al., 2013). However, even when using a minibatch approximation, the KL-divergence term must be calculated at each gradient step, and from it costs ~ *DK*^3^ to evaluate (eq. 45). For large values of *D*, which are expected in high-dimensional neural datasets, the KL-divergence term evaluation makes stochastic gradient descent far too slow.

We solve this problem by representing *W*^⊤^ using a basis 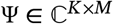, i.e.*W*^⊤^ = Ψ*U*, where 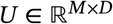. Then, we place a independent normal prior on each entry of *U*. This allows the KL-divergence term to be evaluated with ~ *DM* complexity, since it becomes a KL divergence between two independent normal distributions. The only difficulty is choosing Ψ such that *W* turns out to have the desired Gaussian process distribution when *U* is a independent normal.

It can be shown that the appropriate choice of Ψ is the unitary Fourier basis (see Appendix B). With this basis, the entry *U_id_*, the element in the *d*-th column of *U* corresponding to Fourier frequency *i*, must satisfy two conditions. The first condition is conjugacy, 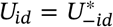, which ensures that *W* is real. The second condition is on the distribution, which must be a zero-mean complex normal with variance

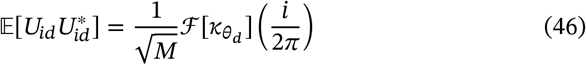

This spectral formulation assumes the stimuli lie on [0, 2*π*], but it can be trivially extended to any domain.

## 5 Results

### 5.1 Evaluation and performance

We benchmarked each decoder by calculating its mean absolute test error on the monkey (Graf et al., 2011), ferret (this paper), and mouse (Stringer et al., 2021) datasets (Figure 3). We chose to estimate the error using five-fold cross-validation repeated ten times, since ten-fold cross-validation produced datasets that had too few examples per class to train some models. Error bars were computed using twice the standard error of the mean of the cross-validation scores 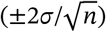. We examined five monkey datasets, one ferret dataset, and three mouse datasets. Figure 3A reports the average scores for each animal; separate scores are reported in Figure S4. Note that the GID decoder has two variants: the standard formulation, which has a quadratic decision boundary, and the formulation described in section 3.1.2, which has a linear decision boundary.

**Figure 3:**
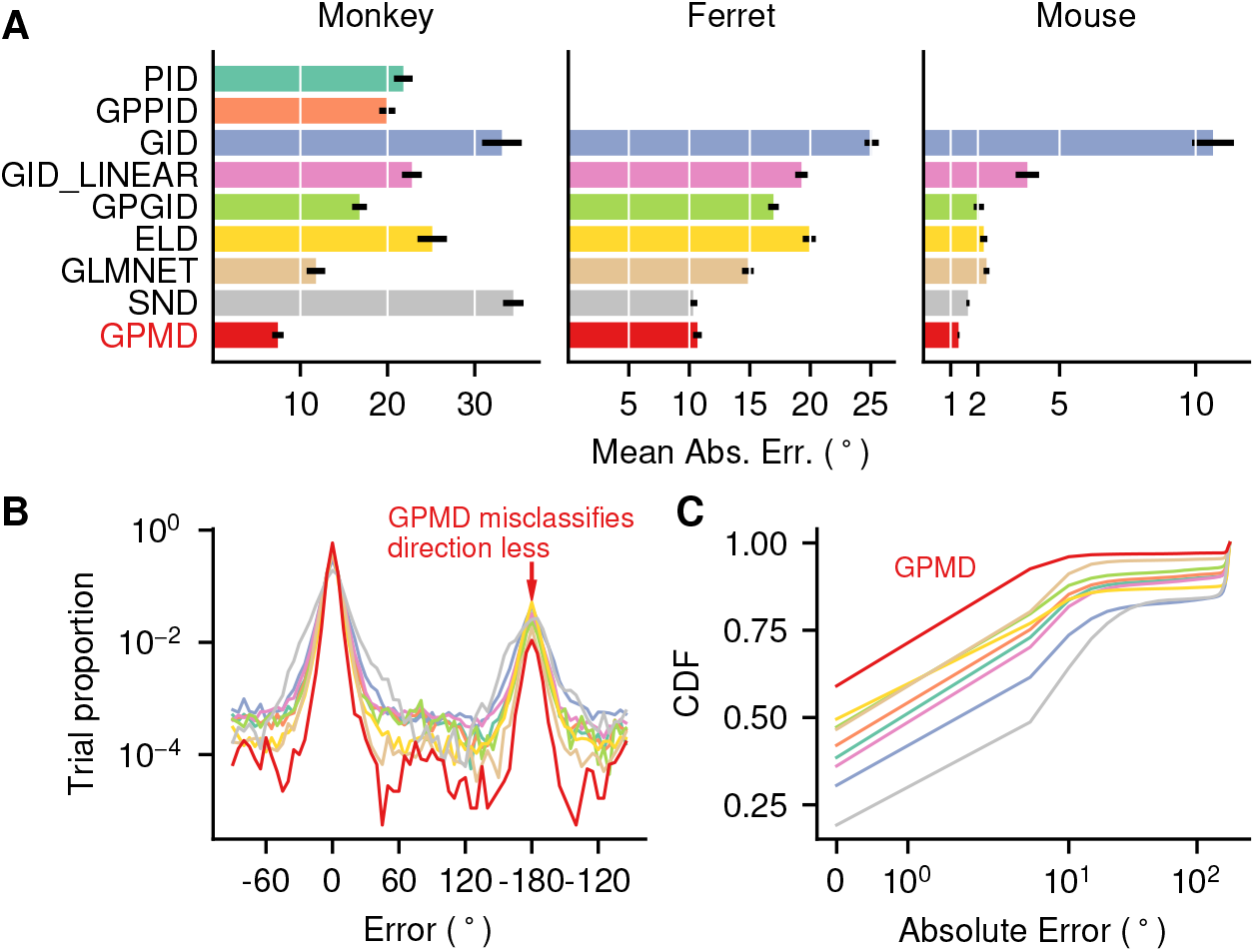
**A**: Test error on monkey (Graf et al., 2011), ferret, and mouse (Stringer et al., 2021) datasets, estimated using five-fold cross-validation repeated ten times. Since there are five monkey and three mouse datasets, we trained the models on each dataset separately and averaged the scores. We did not train the PID and GPPID decoders on the ferret and mouse datasets, which contain real-valued calcium data, because they assume integer features. Error bars are ±2*σ*, where *σ* is the standard error of the mean. **B**: The distribution of errors for each decoder, using the third monkey dataset. The GPMD had more mass concentrated around 0° error (correct classification) and a narrower error distribution than the other decoders. One noteworthy feature of all decoders is a preponderance of errors at 180°, indicating an estimate with the correct orientation but incorrect drift direction. The GPMD nevertheless exhibited this error less frequently than the other decoders. **C**: Empirical CDF of errors on the third monkey dataset. The GPMD classified higher fractions of the dataset at lower errors than the other decoders, demonstrating its superior performance.

The rank ordering of the models remained largely consistent across datasets. In general, the correlation-blind decoders (the PID and GID) performed worse than the correlation-aware decoders, which is consistent with previous decoding studies (Graf et al., 2011; Stringer et al., 2021). Their regularized variants (the GPPID and GPGID) performed better, but still did not match the performance of the best correlation-aware decoders. The GPMD set or matched state-of-the-art performance on all datasets. An important advantage of its Bayesian approach is that hyperparameters are learned automatically, which allowed the GPMD to adapt to the conditions present in different datasets. By contrast, models that set hyperparameters manually exhibited occasional poor performance, for example the SND performance on the monkey datasets.

These results could have been be obscured by the choice of error metric. For example, repeating the same benchmark using “proportion correct” instead of mean absolute error improved the performance of the ELD substantially (see Figure S3), qualitatively replicating the results of Graf et al. (2011). To ensure that our results were not artifacts of the error metric, we used the empirical error cumulative distribution function to characterize each decoder’s errors in more detail (Figure 3B). Good decoders should classify higher fractions of the dataset at lower errors, producing curves that lie up and to the left. We found that the GPMD outperformed or matched all the other decoders on all the datasets (see Figure S5).

Our results show that both regularization and exploiting correlations improved decoding performance substantially. The regularized correlation-blind decoders, the GPPID and GPGID, outperformed their unregularized analogues, the GID and PID. The GLMNET decoder, which is correlation aware, outperformed the correlation-blind GPPID and GPGID. Finally, the SND and GPMD, which are both regularized and correlation-aware, outperformed all other decoders.

The relative impact of these strategies depended on dataset dimensionality. All of the decoders performed better on the higher-dimensional mouse dataset than the mid- and low-dimensional ferret and mouse datasets. This is because as dimensionality increased, the class distributions became further apart, making the problem easier to solve (Figure S2). Increased neural selectivity was correlated with decoding performance (Figure S11). Within each dataset, decoders performed differently depending on how they handled regularization and correlations. For small datasets, such as the monkey dataset with ≈150 neurons, both regularization and exploiting correlations had a substantial effect. For example, adding regularization to the GID (using the GPGID) decreased its mean absolute error by 16.3 degrees, and exploiting correlations (using the GPMD), decreased error by another 9.4 degrees. For high-dimensional datasets where it was easy to overfit, such as the mouse dataset, which was recorded from ~ 20,000 neurons, regularization became the most important strategy. In fact, on the mouse dataset, the regularized correlation-blind GPGID did just as well as some of the correlation-aware decoders.

### 5.2 Scaling with dataset size and dimensionality

To characterize the GPMD’s performance and training times with respect to dataset size and dimensionality, we performed ablation studies on both the number of training examples and the number of neural features (Figure 4).

**Figure 4:**
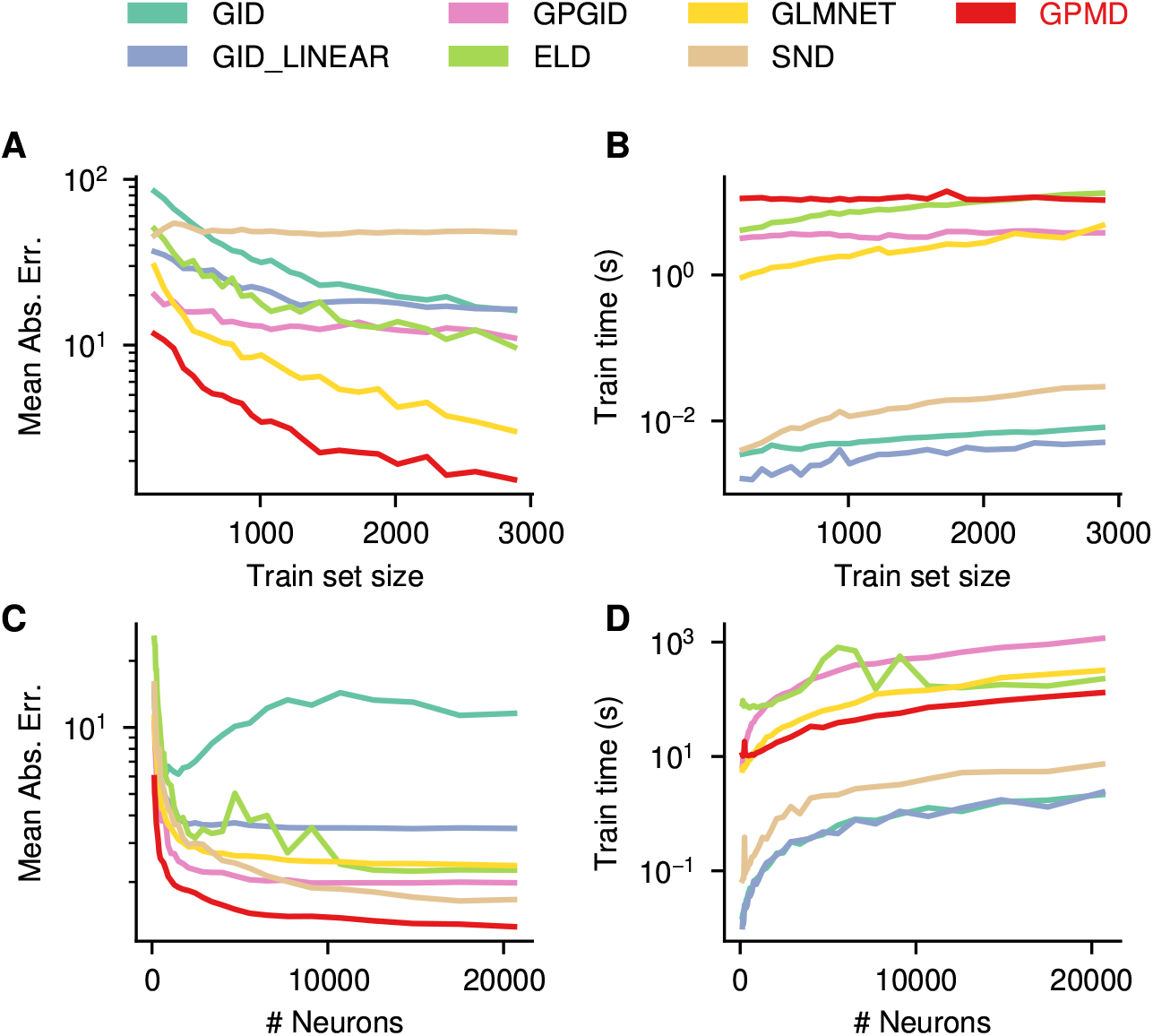
**A**: Cross-validated model performance with varying amounts of training data. We chose random subsets of the third monkey dataset. The GPMD continues to benefit from increasing training data, and does exhibit asymptotic behavior like most of the other models. **B**: Training times for the ablation study in (A). Like all the models, the GPMD shows essentially constant training times. This is because the monkey dataset is small enough that training cost is dominated by constant factors. **C**: Cross-validated model performance with varying amounts of neurons (features). We chose random subsets of the first mouse dataset. The GPMD’s careful regularization avoids undesirable double-descent characteristics while achieving state-of-the-art performance for all feature sizes. **D**: Training times for the ablation study in (C). The GPMD is nearly one order of magnitude faster than logistic regression, a less sophisticated model, and trains in a few minutes on the largest datasets.

The GPMD performed well at all training set sizes (Figure 4A) implying that its prior assumptions encouraging smooth and small decoding weights were well calibrated— that is, strong enough to permit good performance with few training examples, but flexible enough to allow continued learning with many training examples. We believe that the nonparametric nature of the Gaussian Process prior is the key component that allows our model to accommodate the complex structure present in larger neural datasets. Models with stronger assumptions, such as the GPGID, which assumes independence, or the SND, which has many hard-coded parameters, had difficulty learning from increasing numbers of training examples.

The GPMD also performed well with any number of neural features (Figure 4C). Linear decoders with no or poor regularization, such as the GID and ELD, did not exhibit this property; in fact, their performance became worse as the number of neural features increased from the “classical” to the “interpolating” regime, producing a phenomenon known as the “double descent” error curve (Belkin et al., 2019). Properly regularized models such as the GPGID and GPMD did not display this phenomenon and gave accurate performance estimates for all numbers of neural features.

Thanks to the GPMD’s approximate inference, GPU acceleration, and spectral weight representation, it trained quickly, producing fast cross-validated error estimates that exhibited favorable scaling with respect to both observations and neurons (Figures 4B and 4D). For the largest dataset with 20,000 neurons, it took 131 +/- 0.82 seconds to train (roughly 20 minutes of wall-clock time) for a ten-fold cross-validation estimate. By comparison, a performance-tuned GLMNET model took 618 +/- 6.40 seconds to train (roughly 1 hour and 45 minutes of wall-clock time) for the same estimate. Given the training time trends shown in the training-set size ablation (Figure 4BB) and neural feature ablation (Figure 4D) studies, we expect the GPMD to handle even larger datasets without difficulty.

Scaling to large datasets was further enhanced by the GPMD’s automatic dataset preprocessing. Decoding studies, such as Graf et al. (2011), often select only strongly tuned neurons for decoding, since noisy neurons make it easier for models to overfit. Manual selection rules have two disadvantages: first, they may ignore neurons that look noisy but actually carry information, and second, they can require prohibitive amounts of time if human input is needed (e.g., for choosing initialization points for nonlinear curve fitting).

The GPMD’s Bayesian formulation automatically discarded noise neurons by setting their prior amplitudes to zero (Figure 5), a phenomenon known as automatic relevance determination (MacKay, 1992; Neal, 1996). Examples of tuning curves (estimated using the sample average conditional on the stimulus) from automatically discarded and automatically retained neurons are shown in Figure 5. Some of the automatically retained neurons displayed the bimodal “Gaussian bump” structure commonly sought by manual selection rules. Others displayed more complicated tuning patterns that would likely be ignored by a manual selection rule.

**Figure 5:**
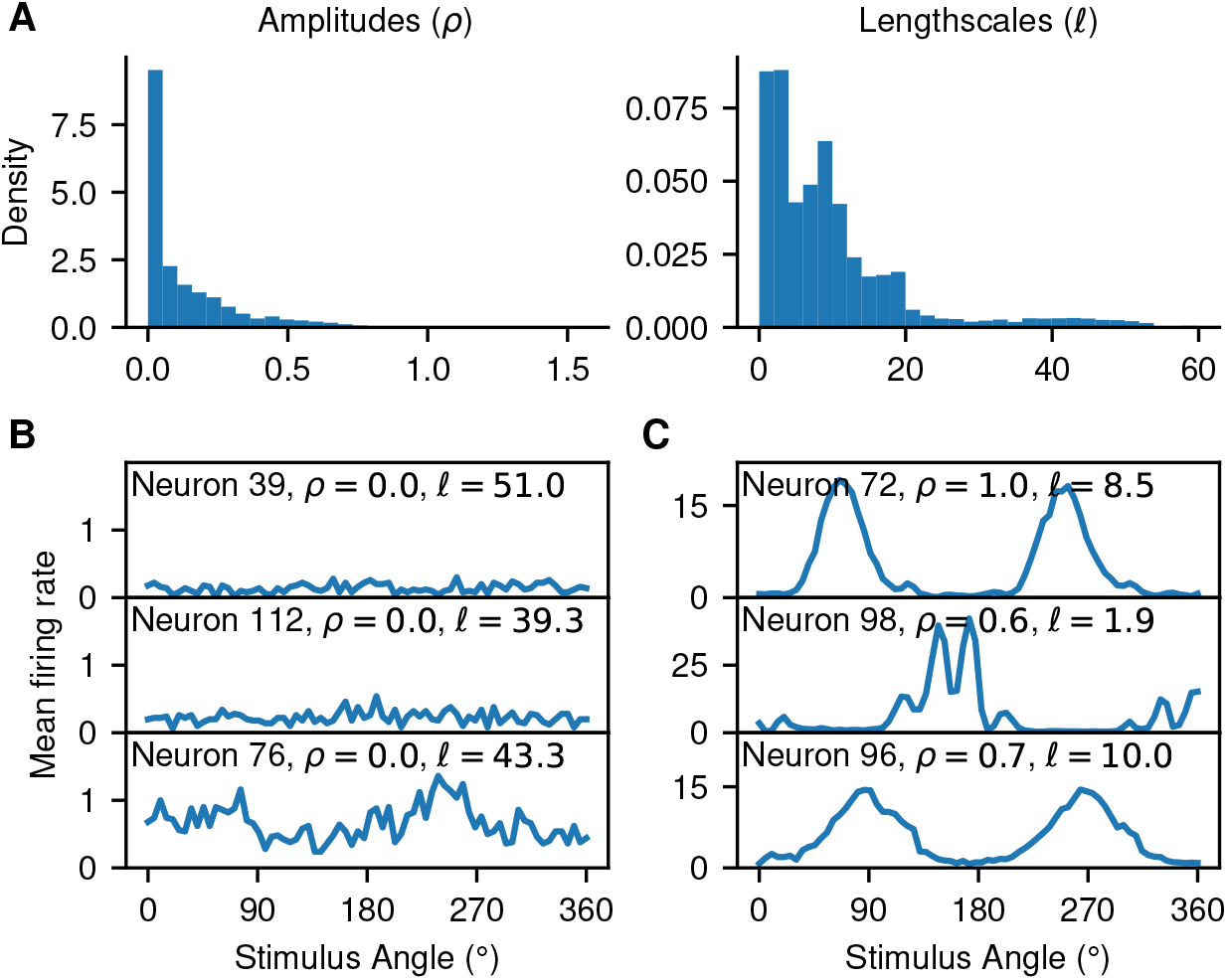
**A**: GPMD hyperparameter distributions on the third monkey dataset. The distribution of amplitudes has a peak near zero, and the distribution of lengthscales has a second mode near 45, indicating an automatic relevance determination (ARD) effect. **B**: Tuning curves for the ARD-eliminated neurons, estimated using the sample average of each neuron’s spike counts conditioned on the stimulus. These neurons have amplitudes near zero and, often, long lengthscales. **C**: Tuning curves for the neurons that were not eliminated. These tuning curves were also estimated using the sample average. These neurons have positive amplitudes and much shorter lengthscales. Some have simple “Gaussian bump” tuning curves, whereas others have more complex tuning characteristics.

Our implementation of the empirical linear decoder (ELD, see Graf et al. [2011]) replicated the original paper’s results only qualitatively, not quantitatively. Our implementation of the ELD did outperform the Poisson Independent Decoder (PID) when using the “proportion correct” error criterion, as in the original paper (see Figure S3). However, it did not achieve the performance reported in the original paper. Because our implementation of the PID, a very simple decoder, also did not match the performance of the PID in Graf et al. (2011), we believe the discrepancy was caused by data preprocessing. We were not able to replicate the data preprocessing steps described in Graf et al. (2011) precisely, since the original code has been lost.

### 5.3 Visualizing the impact of correlations on decoding

As described in section 4, each row of the weight matrix *W* for each decoder forms a vector of decoding weights over orientation for a single neuron. Example decoding weights from a correlation-blind (GPPID for monkey, GPGID for mouse and ferret) and correlation-aware (GPMD) decoder pair are shown in Figure 6A. In general, the GPMD produces smoother weights, and unlike the GPPID/GPGID, it occasionally places large weights on stimuli with low expected response (e.g., ferret neuron 82 and mouse neuron 87). To verify that the GPMD’s decoding weights were usually smoother, we compared the inferred prior lengthscales for each decoder pair (Fig 6B), and found that the GPMD learned longer lengthscales (sign test, *p* < 0.001 for each animal.) We discarded decoding weights with ℓ_2_ norm less than 0.001, since nearly-zero weights indicated a neuron discarded by the learning process, which would have uninformative length scales.

**Figure 6:**
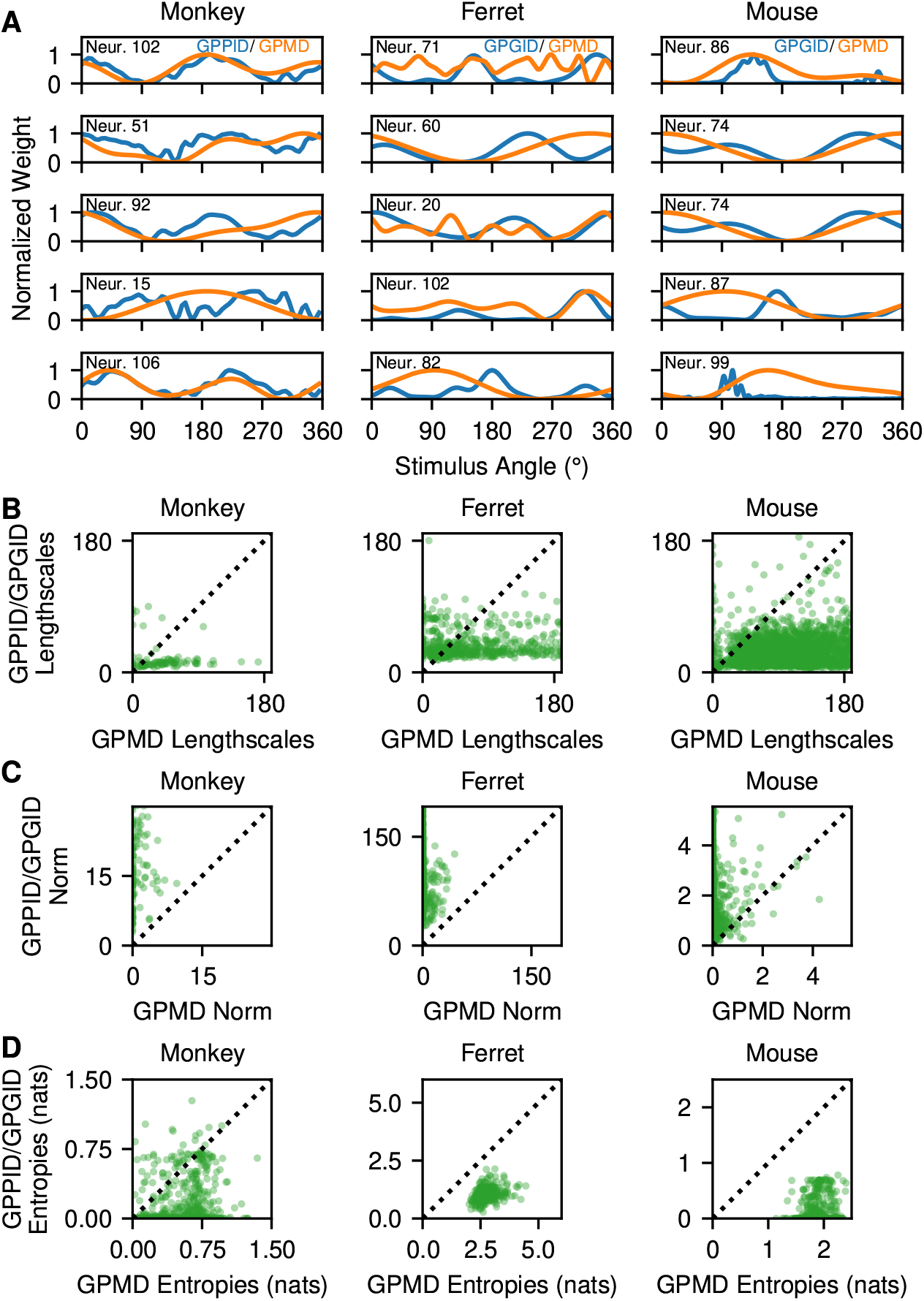
**A**: Inferred decoding weights from naive Bayes (GPPID/GPGID) and GPMD model decoders for five example neurons from monkey, ferret, and mouse datasets. **B**: Comparison of inferred prior lengthscales for each decoder. The GPMD learns longer lengthscales (sign test, *p* < 0.001 for each animal). **C**: Comparison of decoding weight norms for each decoder. The GPMD learns decoding weights with smaller norms (sign test, *p* < 0.001 for each animal). **D**: Comparison of test-set predictive distribution entropies for each decoder. The GPMD produces distributions with larger entropies (sign test, *p* < 0.001 for each animal), indicating that it reports more uncertainty.

In nearly all cases, the GPPID/GPGID decoders learned decoding weights with larger ℓ_2_ norm than the GPMD (Figure 6C, sign test *p* < 0.001 for each animal). Since the norm of the tuning curve is directly related to the predicted probability for each class (Equation 1), this indicates that the GPPID/GPGID are more confident in their predictions than the GPMD. To directly measure this, we chose a test set for each dataset and compared the predictive distribution entropy for both the GPPID/GPGID and GPMD decoders (Figure 6D). We found that the GPMD produced predictive distributions with higher entropy (sign test, *p* < 0.001 for all animals), indicating that it produces more uncertain predictive distributions. This confirms the patterns seen in Figure 6C.

To further characterize the effects of correlations on decoding performance, we visualized population responses projected onto 2D subspaces defined by the decoding weights of correlation-blind (GPPID) and correlation-aware (GPMD) decoding models (Figure 7A). To reduce the 147-dimensional response vectors obtained from a population of 147 monkey V1 neurons, we first selected a model to visualize, and two stimulus classes *i* and *j*. Then, we formed a two-dimensional basis by orthogonalizing the decoding weight vectors *W_i*_* and *W_j*_*, producing a basis matrix of orthogonal unit vectors, *S*. We plotted the data for classes *i* and *j*, and the model separatrix by projecting onto this basis. The model separatrix is given by the solution to the equation **x**^⊤^**u** + *c* = 0, where **u** = *S*(*W_j*_* – *W_i*_*) and the constant offset *c* is **b**_*j*_ – **b**_*i*_. We estimated the approximate separatrix from the model that did not produce the basis by projecting its normal vector into the subspace defined by *B* and setting the constant offset to be −(mean(*X_i_***u**) + mean(*X*_j_**u**)/2, where *X_i_* and *X_j_* are the data for classes *i* and *j*.

**Figure 7:**
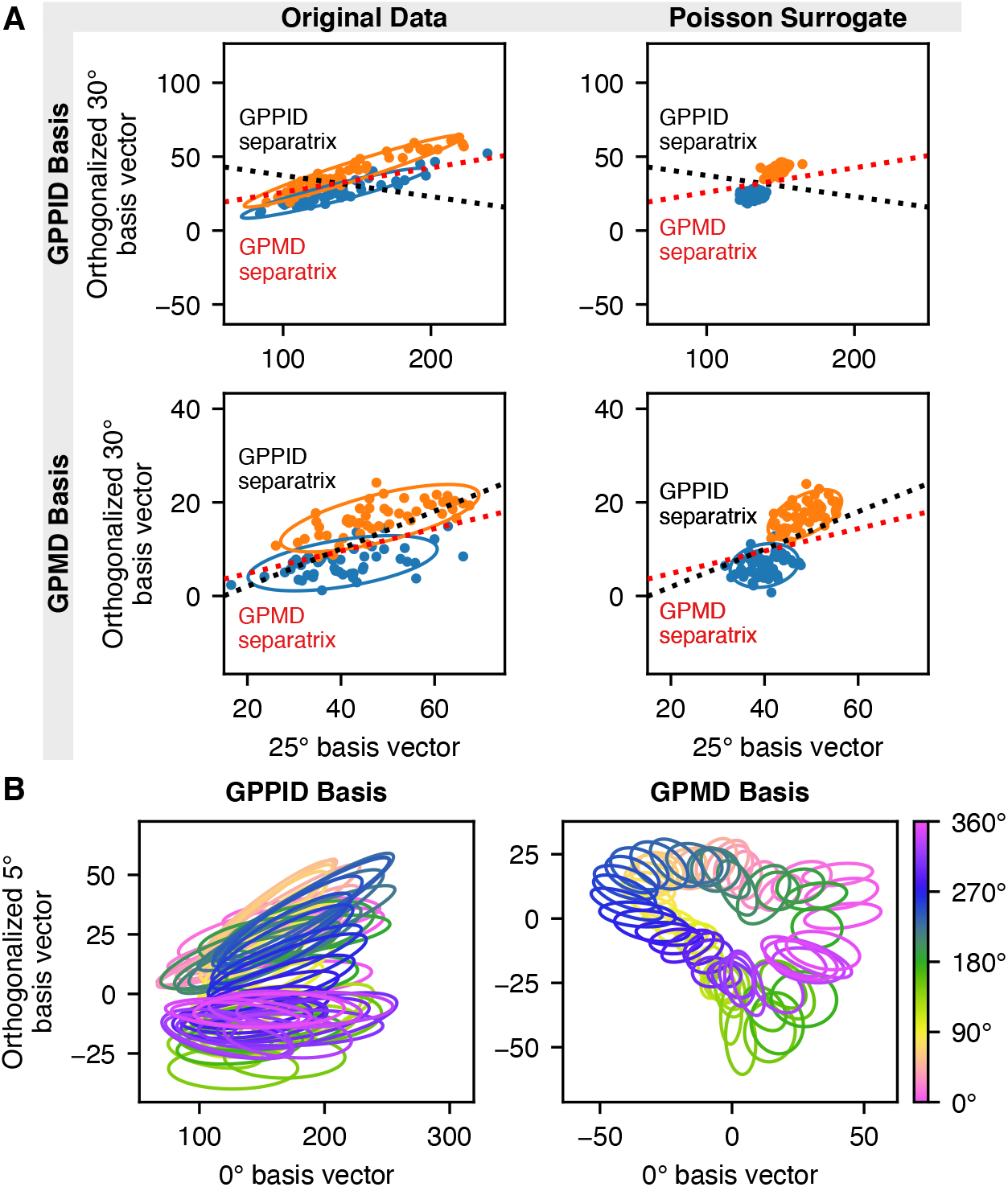
**A**: To investigate the impact of correlations on decoding performance, we projected the 25° and 30° data classes from the third macaque dataset to two dimensions using an orthogonalized bases derived from the 25° and 30° decoding vectors from the correlation-blind GPPID and correlation-aware GPMD. The data displayed significant correlations in the GPPID basis, making the GPPID separatrix a poor decision boundary. The GPPID separatrix only performed well when correlations were removed from the dataset using a Poisson surrogate model. In the GPMD basis, the difference between the correlated and Poisson surrogate datasets was much less pronounced, indicating that the GPMD’s projection decorrelated the data somewhat. **B**: Class correlation ellipses plotted using the GPPID and GPMD bases derived from zero- and five-degree decoding vectors. The GPMD basis produced far superior class separation.

We carried out this procedure for both the GPPID and GPMD models, since each two-dimensional basis could exactly represent only the separatrix from its source weight matrix. Figure 7A shows our visualization for the 25°/30° class pair, and we included four additional class pairs in Figure S9 (all class pairs exhibited a similar pattern).

We first wished to determine whether the data deviated significantly from the independent Poisson model assumed by the GPPID. To do this, we generated a simulated dataset from a conditionally independent Poisson model with means given by the tuning curves of the GPPID model (referred to in the figure as the “Poisson surrogate” dataset). Compared to the real data, the Poisson surrogate data exhibit much smaller variance and tilt relative to the basis vectors, showing that the correlations in the real data can significantly affect decoding. However, in the GPMD basis, the differences between the real and surrogate datasets are much less pronounced, implying that the GPMD’s weight matrix identifies a linear projection that decorrelates and reduces the variance of the projected data. We found the same results using a decorrelated datset created by shuffling neural responses across trials (Figure S8).

Next, we plotted the class separatrices along with the data. The separatrix given by the GPPID successfully separated the Poisson surrogate data, but failed to separate the real dataset because of correlation-induced distortions. However, as expected, the GPMD separatrix successfully took the data’s correlations into account.

To visualize how the entire set of 72 classes related to each other, we plotted each class’s correlation ellipse on the basis given by the zero- and five-degree basis vectors from each model (Figure 7B). The selection of the basis vectors was arbitrary, see Figure S10 for other choices. The GPPID’s basis did a poor job of separating the classes, but the GPMD’s basis separated them fairly well. In the GPMD’s basis, the ellipses from classes 180 degrees apart appear in nearly identical locations, confirming that that the GPMD identified grating angles more precisely than grating drift direction, a phenomenon previously observed in our performance benchmarks (Figure 3C).

## 6 Discussion

Linear decoders provide a powerful tool for characterizing the information content of neural population responses. While all linear decoders share a common mathematical form, differences in fitting methods—namely, in regularization and the incorporation of noise correlations—can lead to large differences in performance.

Previous studies, such as Graf et al. (2011) and Stringer et al. (2021), have used correlation-blind and correlation-aware linear decoders to investigate the significance of correlations on decoding performance. In all cases, they found that correlation-aware decoders outperform correlation-blind decoders. However, the performance differences could have been due to the lack of regularization in the correlation-blind decoders, as maximum likelihood weight estimates in the sample-poor data regimes typically studied are often subject to overfitting.

In this paper, we presented a suite of new decoders that use Gaussian processes for regularization. Two of these methods correspond to naïve Bayes decoders under Poisson and Gaussian noise models, respectively, which do not incorporate correlations. The third method, the GPMD, corresponds to a decoding model with a GP prior over its weights that does take account of noise correlations. We found that the GPMD matched or outperformed all other decoders on datasets from monkey, ferret, and mouse visual cortex. Furthermore, we showed that the GPMD scales to the very largest datasets using a combination of approximate Bayesian inference, spectral methods, and GPU acceleration.

Our results show that the performance gap between correlation-blind and correlation-aware decoders is fundamental, and not just a result of regularization. The correlation-aware decoders consistently outperformed even the regularized correlation-blind decoders, though regularization did narrow the gap significantly (as shown in Figure 3). This confirmed the results of previous studies, and we conclude that exploiting neural correlations can significantly improve decoding performance (though this result may have some dependence on whether or not the animal is anesthetized, see Yates et al., 2020). Visualizations of the decoding separatrices produced by each decoder indicate that the real datasets differ significantly from the assumptions made by correlation-blind decoders. The correlation-aware decoders discovered low-dimensional subspaces that decorrelated the responses, making the transformed data match independence assumptions more closely.

One major caveat to our results is that they cannot tell us whether the brain itself uses decoders that are correlation-blind or correlation-aware. Animal behavior may rely on a different populations of neurons than the ones recorded in our experimental datasets. Animals might achieve similar performance to the GPMD by reading out a similar neural population optimally or a larger neural population sub-optimally (Chen et al., 2006; Jacobs et al., 2009; Morais et al., 2022). As in previous studies (Stringer et al., 2021), we found that in the largest recordings from mice, the decoding performance (1.3°) substantially exceeded current estimates of discrimination ability of the animal (9°; Lyamzin et al. (2021)). This suggests that the animal’s behavioral performance may be primarily limited by the decoding strategies implemented by downstream regions, and not by the information present in V1 itself (Beck et al., 2012).

The prior assumption of smoothness governs the applicability of the GPMD and other GP-regularized decoders we developed to other brain regions and tasks. If the neural population of interest has smoothly varying tuning curves and the task can be solved using a linear decision boundary, the GPMD will perform well. Although our decoding task used angular stimuli and a periodic distance measure, the GPMD could be trivially extended to stimuli that vary smoothly on an interval of the real line by using the ordinary Euclidean distance. Beyond smoothness, we have made no assumptions about the population of neurons to be decoded, and the GPMD naturally “prunes” uninformative neurons (e.g., neurons untuned to the stimulus) by shrinking their decoding weights to zero.

In the future, similar decoders could be used to empirically characterize how the peak and high-slope regions of tuning curves contribute to optimal decoding strategies in real neural populations (for theoretical studies, see Butts and Goldman [2006] and Yarrow et al. [2012]). We also believe that the performance of the decoders could be further improved. In particular, the performance of the correlation-aware decoder could likely be improved by modeling additional structure in the neural dataset—for example, the differences in response characteristics between excitatory and inhibitory neurons, or temporal structure (which would likely improve the classification of direction). The correlation-blind decoder could also be improved by using a generative model that can handle non-Poisson characteristics of neural data such as overdispersion (Charles et al., 2018; Gao et al., 2015; Goris et al., 2014; Gur et al., 1997; Stevenson, 2016). We are optimistic that future work will use and extend the decoding methods we have introduced to obtain new insights into the structure of neural codes.

## Acknowledgements

This work was supported by grants from the Simons Collaboration on the Global Brain(SCGB AWD543027), the NIH BRAIN initiative (NS104899 and R01EB026946), and a U19 NIH-NINDS BRAIN Initiative Award (5U19NS104648). Jacob L. Yates is supported by the NIH (K99EY032179). Benjamin Scholl is supported by the NIH (K99EY031137) and thanks the Max Planck Society and Max Planck Florida Institute for their generous support. We thank A. B. A. Graf, Al Kohn, M. Jazayeri, and J. A. Movshon for providing the primate datasets; C. Stringer, M. Michaelos, and M. Pachitariu for providing the publicly available mouse datasets; and Kenneth La-timer for helpful comments on the manuscript.

## A Linearity of the PID and GID decoders

Under appropriate assumptions, both the GID and the PID decoders have linear decision boundaries. To derive both decision boundaries at the same time, let us consider the more general case of a naïve Bayes decoder as described in section 3.1.1, but with an exponential family likelihood (Bishop, 2006). That is, the likelihood of the *d*th element of the feature vector *x_d_* can be written as

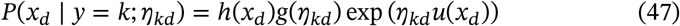

where *η* is the natural parameter and the sufficient statistic 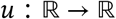 is a function of *x_d_*.

With this likelihood in mind, we can begin solving for the decision boundary. Our class prediction 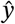 for a given example **x** is

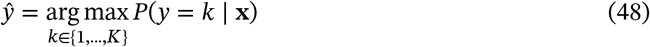

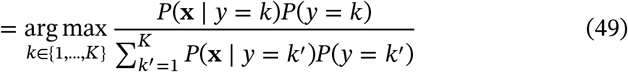

If we assume that the prior probabilities are constant, i.e. *P*(*y* = *k*) = *P*(*y* = *k*′) for every *k*, *k*′ ∈ {1, …, *K*}, then this simplifies to

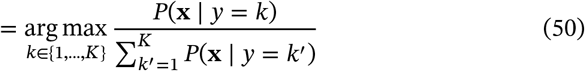

Introducing a log under the argmax, dropping terms that don’t depend on *k*, and substituting 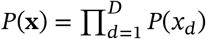, we simplify further:

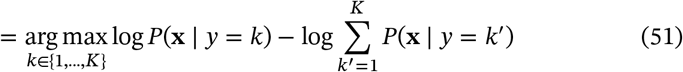

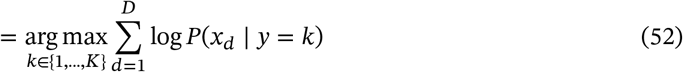

Writing out the exponential family form, we have

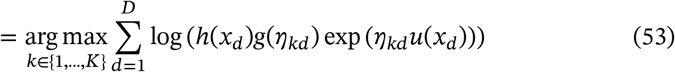

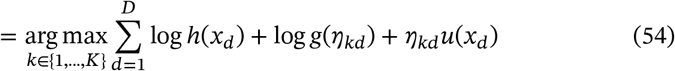

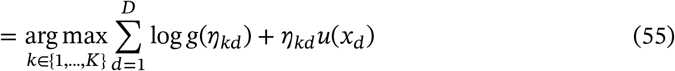

This will simplify to the form of a linear decoder as long as the sufficient statistic of the exponential distribution *u*(*x_d_*) of the form *u*(*x_d_*) = *αx_d_* where *α* is a scalar. In that case, the entries of the weight matrix are given by the natural parameters

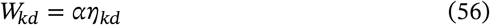

and the entries of the intercept vector **b** are given by

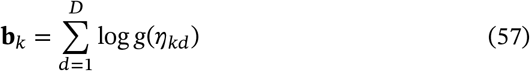

Using these definitions, we can write

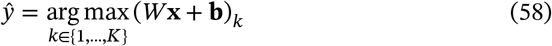

which is the form of a linear decoder.

In the case of the Poisson Independent Decoder, the sufficient statistic is the identity function, the natural parameter of the Poisson distribution is given by *η_kd_* = log *λ_kd_* and *g*(*η_kd_*) = *e*^−*λ_kd_*^. Thus, for the PID,

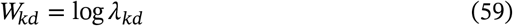

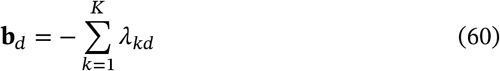

The case of the Gaussian Independent Decoder is slightly more complicated, since the Gaussian sufficient statistic only takes the proper form if the variance *σ*^2^ can be incorporated into *h*(*x_d_*), a term we dropped from the argmax. For this dropping to be valid, we must constrain *σ_kd_* = *σ*_*k*′*d*_ for all *k*, *k*′ ∈ {1, …, *K*}. If this is true, then the sufficient statistic is *u*(*x_d_*) = *x_d_*/*σ_d_*, the natural parameter is given by *η_kd_* = *μ_kd_*/*σ_d_*, and 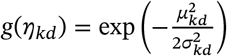. Thus, for the GID,

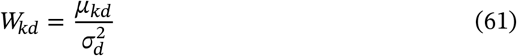

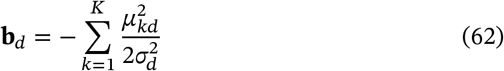

## B Spectral GP regression

### B.1 With Gaussian noise

In this section we demonstrate how to solve a 1-D GP regression problem in the spectral domain, following the general approach of Paciorek (2007), Royle and Wikle (2005), and Wikle (2002). Consider a regression dataset 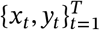 with 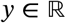 and, without loss of generality, *x* ∈ [−*π*, *π*]. Note that we do not require the *x* values to lie on a grid. We can concatenate the training examples and labels into vectors as follows: **x** = (*x*_1_,.,*x*_*t*_)^⊤^ and **y** = (*y*_1_,…,*y_t_*)^⊤^.

We assume that the values of **y** are noisy observations of a zero-mean Gaussian process *z* where **z_*i*_** = *z*(**x_*i*_**), i.e. ***y_i_*** = **z_i_** + *ϵ* where 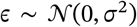. Our probability model can be written:

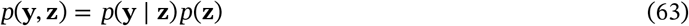

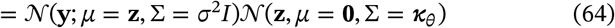

Here ***κ**_θ_* is the kernel matrix between **x** and **x**. If *κ_θ_*(·) is the stationary kernel function with hyperparameters *θ*, then (***κ**_θ_*)_*ij*_ = *κ_θ_*(**x***_i_* – **x**_*j*_). To infer the GP hyperparameters using type-II maximum likelihood, we wish to maximize the log evidence given by

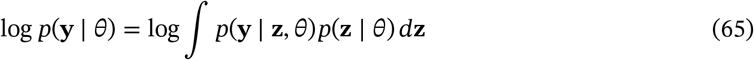

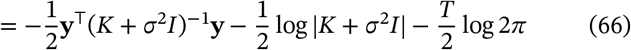

However, each evaluation of this expression has cost ~ *n*^3^, which is intractable for large *n*. Our goal is to decorrelate **z** so that the prior covariance becomes diagonal, dropping the cost to ~ *n*.

To achieve this we will represent the GP *z*(*x*) using a Fourier series:

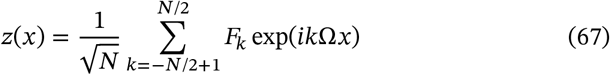

where Ω = 2*π*/*N* and the *F_k_* are the Fourier series coefficients. Our goal is to find the distribution of *F_k_* such that z is a real zero-mean Gaussian with Cov[*z*(*x*), *z*(*x*′)] = *κ_θ_*(*x,x*′).

To ensure that *z*(*x*) is real, we require 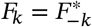. This requirement can be verified by expanding equation 67 in terms of sines and cosines. To ensure that *z*(*x*) is Gaussian, we require that the *F_k_* are independent Gaussian random variables, which implies that they are jointly Gaussian. Since *z*(*x*) is a linear combination of the *F_k_*s, this implies it is also Gaussian. To ensure that *z*(*x*) is zero mean, we require that each *F_k_* is zero mean. Because expectation is a linear operator, this ensures that *z*(*x*) is also zero mean.

The trickiest task is finding the variance of the *F_k_*s that induces the proper GP distribution on *z*. We can construct an equation to solve for it as follows: given a lag *τ* = *s* – *t* between two *x* values, we have the covariance

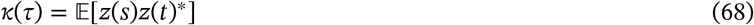

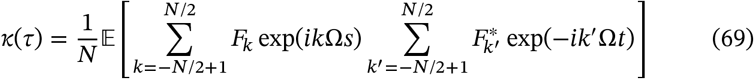

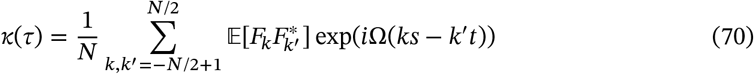

Because the Fourier coefficients are independent, we have 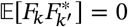 for all *k* ≠ *k*′.

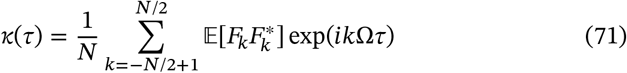

This is just a Fourier series. Thus, we can use the Fourier coefficient formula to invert the equation and solve for 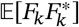:

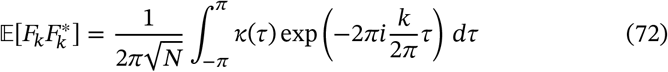

At this point, we are done. Note that the resulting kernel is positive definite, since equation 71 evaluates to a linear combination of cosine kernels, which are positive definite (Murphy, 2012). Many implementations use the Fourier transform of *κ* rather than the Fourier coefficient expression given above. The equivalence can be derived by extending the bounds of integration to [−∞, ∞]. This is a reasonable approximation as long as *κ*(*τ*) is close to zero outside [−*π*, *π*]—which is true for the RBF and Matern kernels as long as the lengthscale is short. Extending the bounds of integration, we have

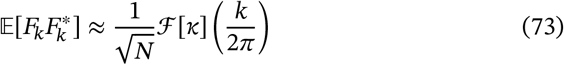

For notational simplicity, let **w** be the vector of frequency-domain coefficients and **s** be the vector of associated covariances. Then the log evidence we wish to maximize is

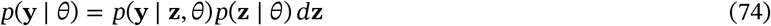

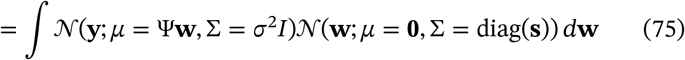

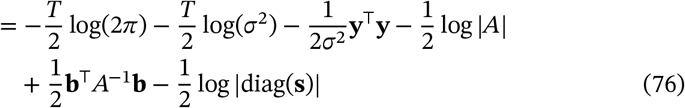

where 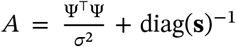 and 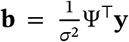. If Ψ is unitary, which is true if **x** lies on a grid, then *A* will be diagonal, which simplifies the gradient and Hessian calculations somewhat.

### B.2 With Poisson noise

Consider the same regression problem as in §B.1, but with Poisson observation noise. We wish to fit the hyperparameters by maximizing the log evidence

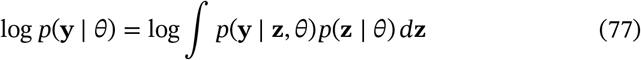

Since 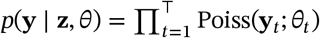 the integral is not analytically tractable.

Define 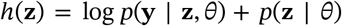 and its argmax as **z**\*. To use the Laplace Approximation (Azevedo-Filho & Shachter, 1994) to approximate the integral, we will start with solving for the evidence using Bayes’ rule:

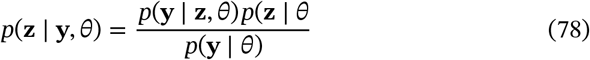

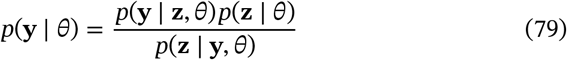

Now, define *H* to be the Hessian matrix of *h*(**z**) and approximate the posterior *p*(**z** | **y**, *θ*) as a Gaussian, using the Laplace approximation:

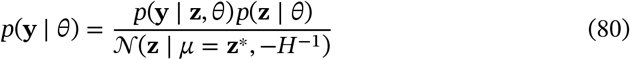

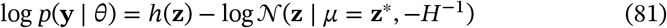

Because log *p*(**y** | *θ*) does not depend on **z**, we will evaluate the right-hand side at **z** = **z**\* to simplify the normal distribution:

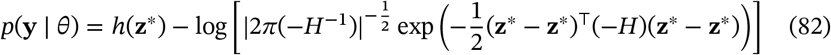

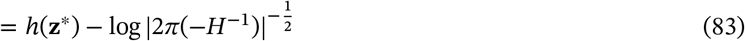

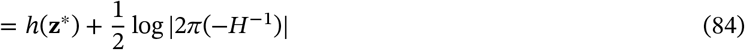

Let *N* be the dimension of **z**.

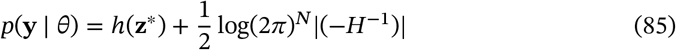

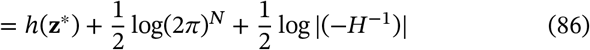

Using the identity log |*A*^−1^1 = log 1/*A* = – log *A*, we have

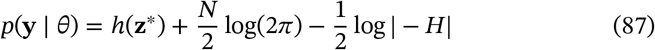

Since evaluating this quantity requires finding **z**\* via an optimization procedure, it is difficult to maximize it using a derivative-based optimization algorithm, an issue pointed out by Rasmussen and Williams (2006). We use a derivative-free technique, the Nelder-Mead algorithm (Nelder & Mead, 1965).

## C Dataset and preprocessing details

For each of the five monkey datasets provided by Graf et al. (2011), we chose the feature ({**x**}) exactly as in Graf et al. (2011). The stimuli grating angles were selected from a five-degree grid, so to get *y_t_* we simply mapped the angles {0, 5, 10, …, 360} to the integers {0, 1, 2, …, 72}.

Unlike Graf et al., we did not drop noisy neurons from the dataset, since we found it made little to no difference in decoding accuracy (see Figure S6).

For the three mouse datasets, we chose the feature vectors 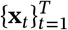 exactly as in Stringer et al. (2021). For the class values 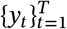, we binned the stimulus angles using 2-degree bins and used the bin index as the class label.

For the ferret dataset, all procedures were performed according to NIH guidelines and approved by the Institutional Animal Care and Use Committee at Max Planck Florida Institute for Neuroscience. Surgical procedures and acute preparations were performed as described in Scholl et al. (2017). To preform calcium imaging of cellular populations, AAV1.Syn.GCaMP6s (UPenn) was injected at multiple depths (total volume 500 nL). Visual stimuli were generated using Psychopy (Peirce, 2007). The monitor was placed 25 cm from the animal, centered in the receptive field locations for the cells of interested. Square-wave drifting gratings (0.10 cycles per degree spatial frequency, 4Hz temporal frequency) were presented at 2 degree increments across the full range of directions (1 second duration, 1 second ISI, 11 trials). Two photon imaging was performed on a Bergamo II microscope (Thorlabs) running Scanimage (Pologruto et al., 2003) (Vidrio Technologies) with 940nm dispersion-compensated excitation provided by an Insight DS+ (Spectraphysics). Power after the objective was 40 mW. Images were collected at 30 Hz using bidirectional scanning with 512×512 pixel resolution. The full field of view was 1 × 1 mm. Raw images were corrected for in-plane motion via a non-rigid motion correction algorithm (Pnevmatikakis & Giovannucci, 2017). Regions of interest were drawn in ImageJ. Mean pixel values for ROIs were computed over the imaging time series and imported into MATLAB (Hiner et al., 2017). Δ*F*/*F_o_* was computed by computing *F_o_* with time-averaged median or percentile filter. Δ*F*/*F_o_* traces were synchronized to stimulus triggers sent from Psychopy and collected by Spike2. Response amplitudes for each stimulus on each trial was calculated as the sum of the Fourier mean and modulation (*F*_0_ + *F*_1_). These values for each neuron were used to generate the feature ({**x**}) vectors. Class values (y) were the stimulus angles presented (at 2-degree increments), using the bin index as the class label.

## D Supplementary figures

**Figure S1:**
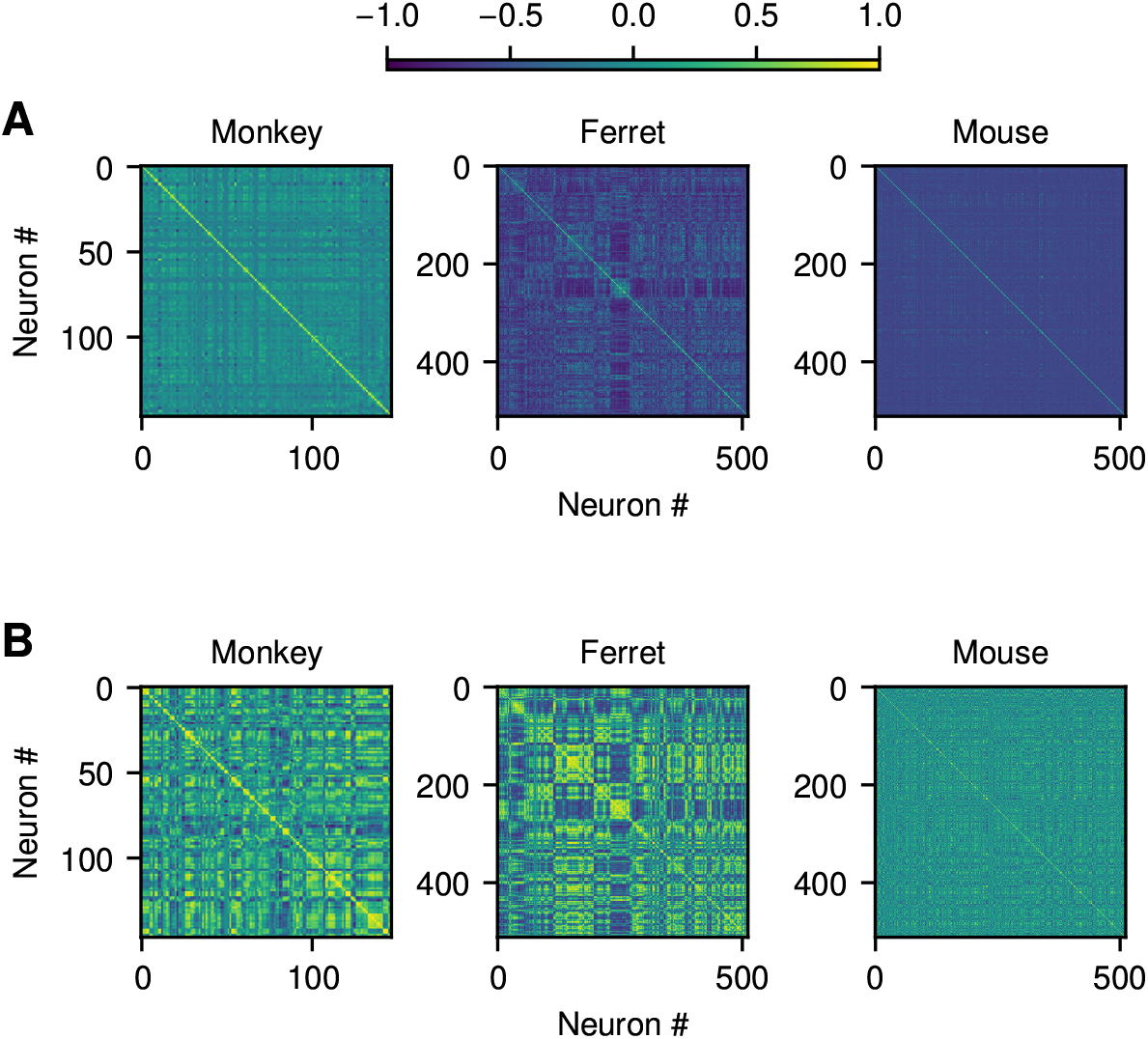
Correlations present in the data. **A:** Noise correlations calculated by taking the correlation coefficient of the neural responses, minus the mean response conditional on the stimulus. The mean responses were estimated using the GPPID or GPGID models, as appropriate, since these were less noisy than the tuning curves estimated using the sample mean. We restricted the ferret and mouse datasets to the first 512 neurons so that the patterns would be large enough to be visible. **B:** Signal correlations, calculated by computing the correlation coefficients of the neural tuning curves. In this case, we estimated the neural tuning curves using the GPPID/GPGID models. As before, we restricted the ferret and mouse datasets to the first 512 neurons.

**Figure S2:**
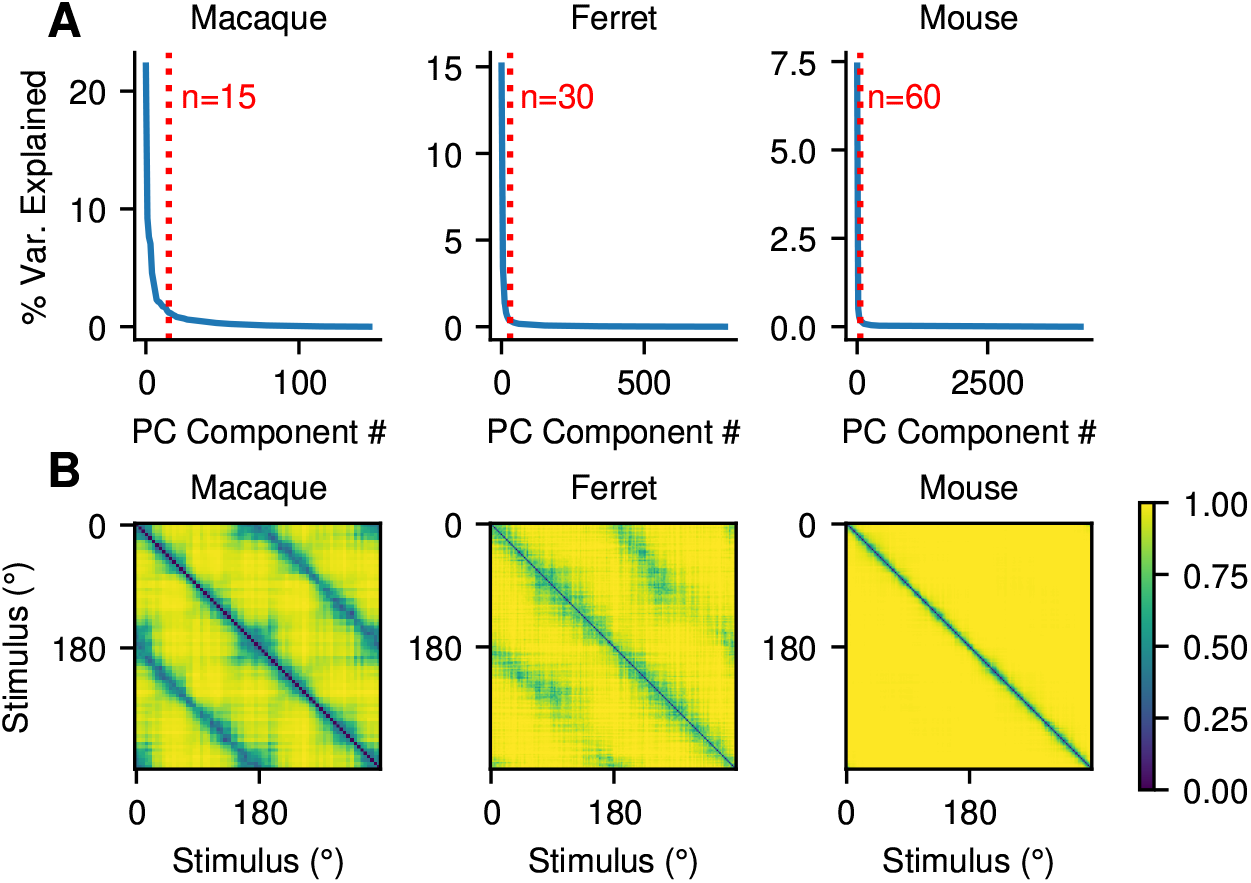
A visualization of the distance between class distributions in each dataset. **A:** To prevent numerical instability in our calculations, we first reduced the dataset dimension using PCA. The number of components chosen is marked in red. **B:** Distance between class distributions, estimated using the analytic Hellinger distance bewteen Gaussian distributions fit empirically to each class distribution. The Hellinger distance is a symmetric distance measure that takes the value zero when the distributions are maximally similar and one when the distributions are maximally dissimilar. Nearby class distributions are similar to each other, and to distributions 180° away. As dimensionality increases, the class distributions become more and more separated.

**Figure S3:**
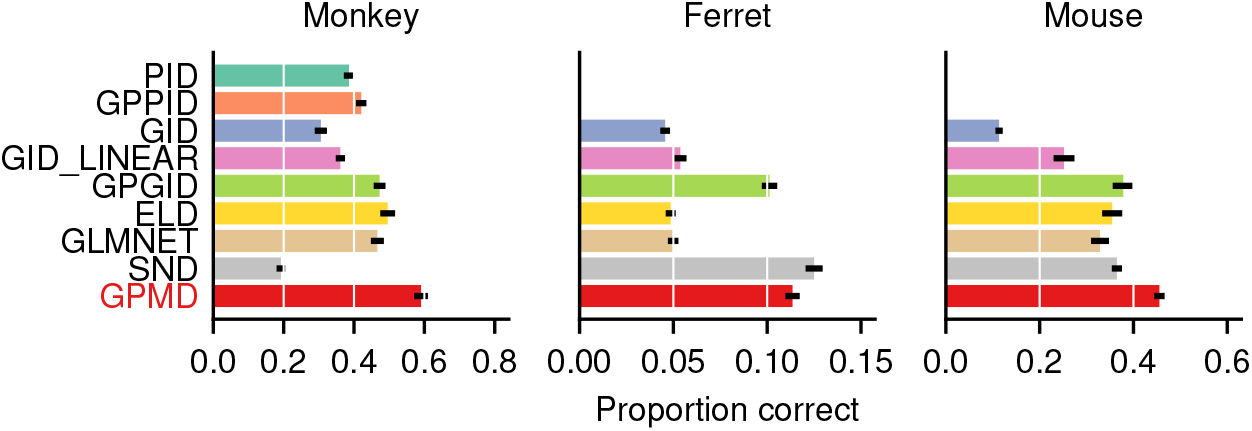
The same benchmark as in Figure 3, but calculated using proportion correct (i.e. proportion with 0° error) instead of mean absolute error. This is the same criterion as in Graf et al., 2011. Using it, we qualitatively replicate the results of Graf et al.

**Figure S4:**
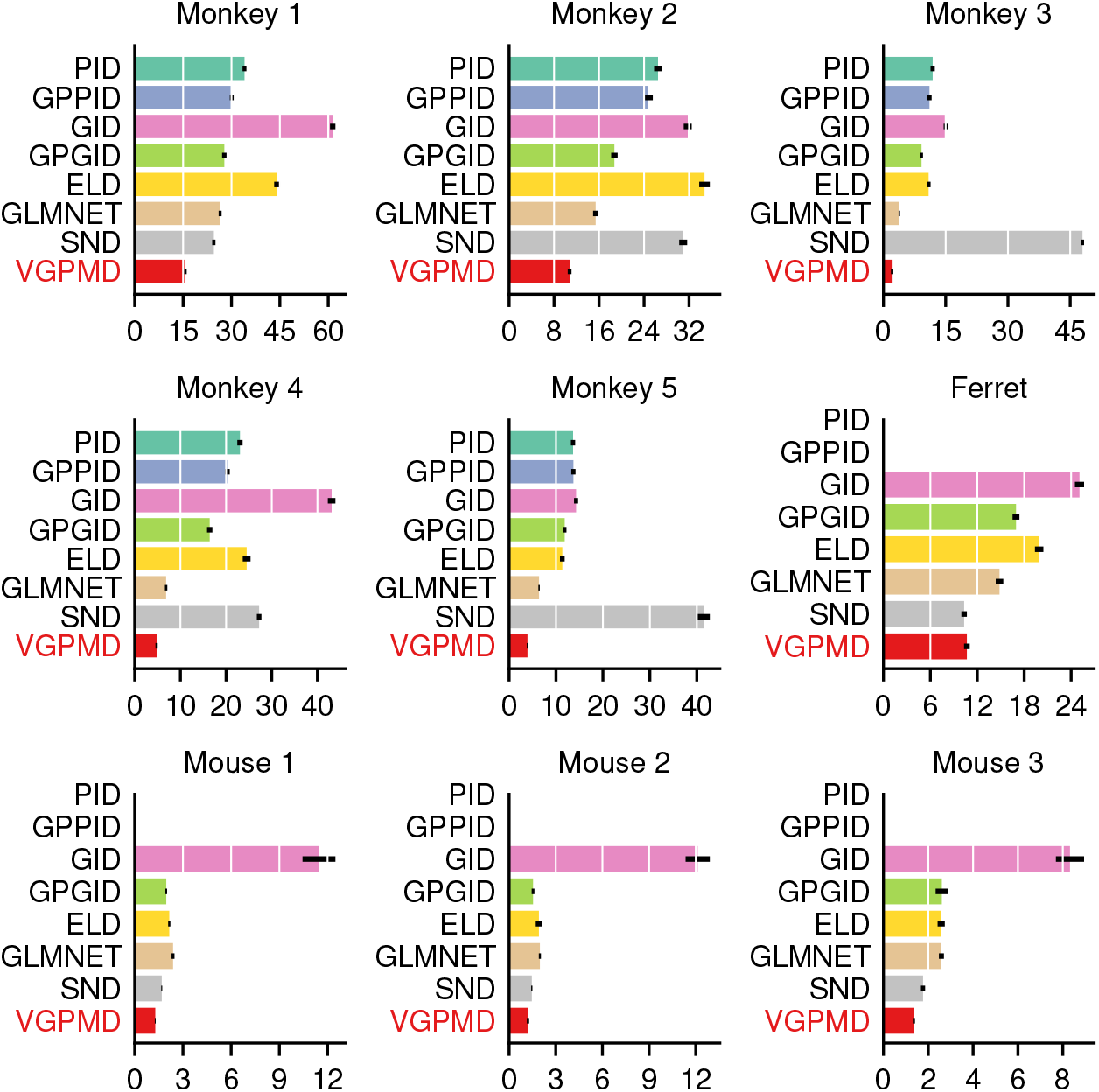
The same benchmark as in Figure 3, but without averaging across animals.

**Figure S5:**
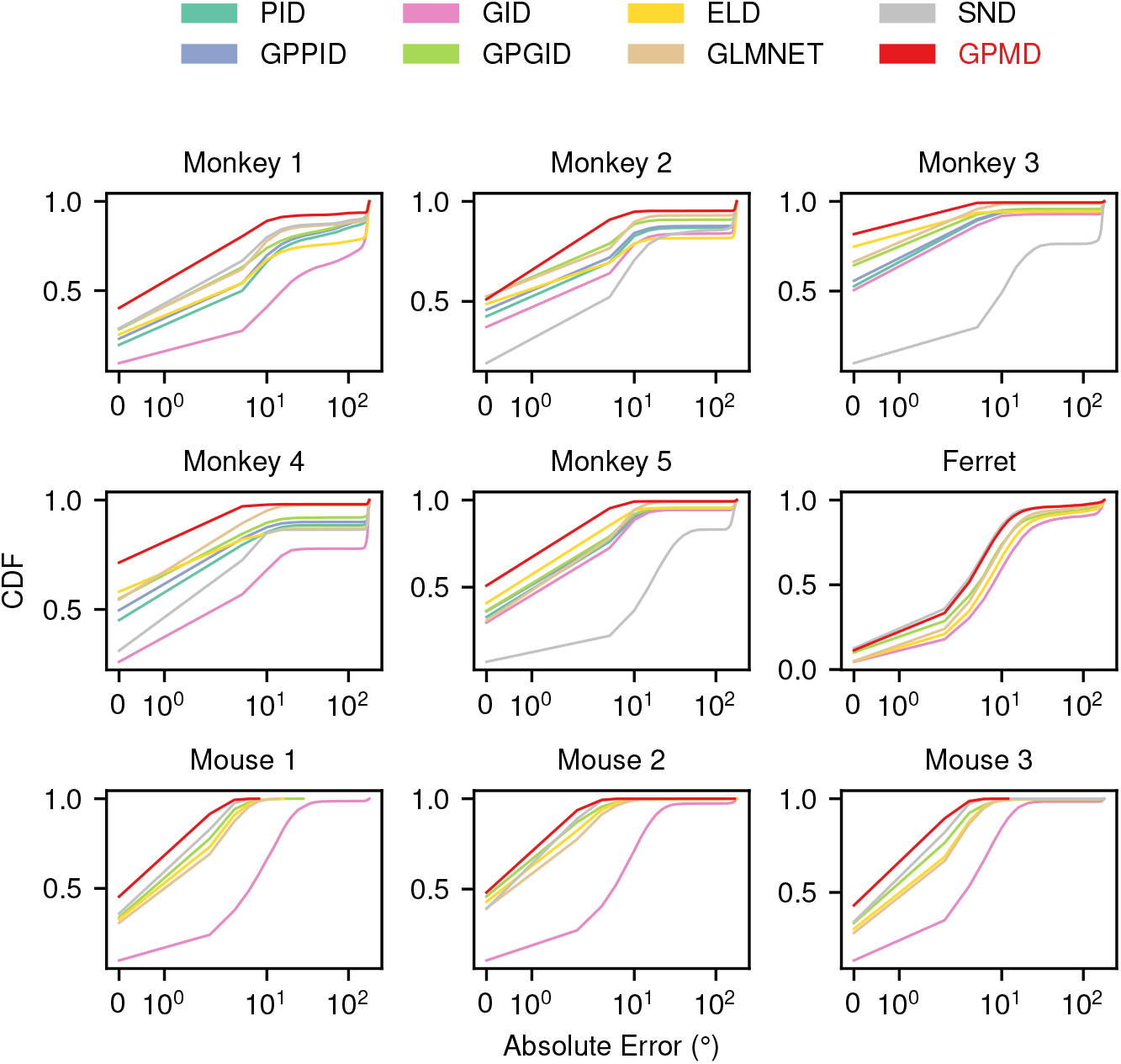
The same benchmark as in Figure 3B, but without averaging across animals.

**Figure S6:**
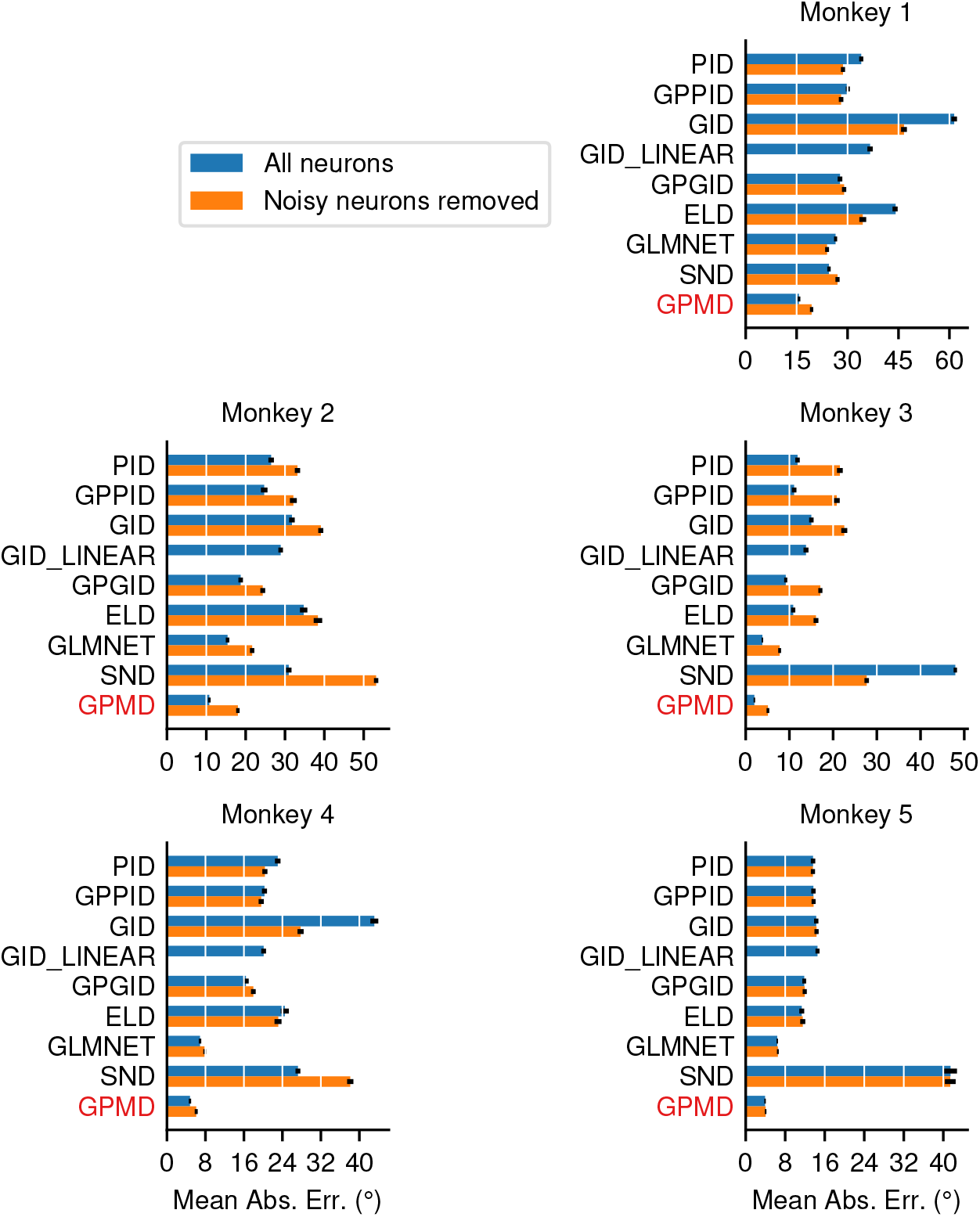
A comparison of model performance on the monkey datasets from Graf et al. (2011), both with and without noisy neurons included. Noisy neurons were dropped using the procedure detailed in the supplementary information of Graf et al. (2011). We thought that this filtering step would make the models perform better, matching the performance reported by Graf et al. (2011), but it did not. We suspect that the issue lies in the initialization of our nonlinear curve fitting code, but we were not able to compare our implementation with the original, since the original code has been lost.

**Figure S7:**
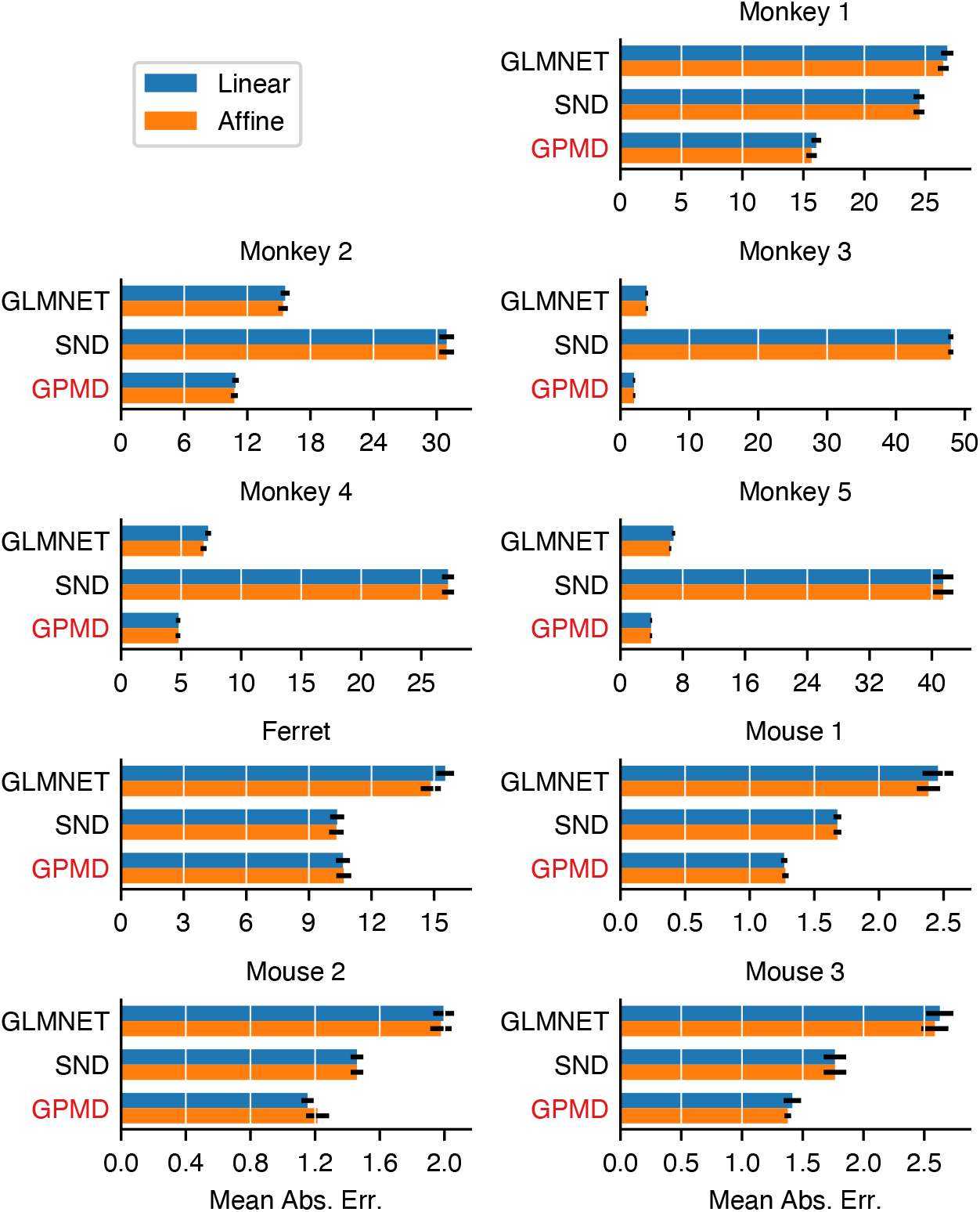
A comparison of model performance using linear (*y* = **w**^⊤^**x**) and affine (*y* = **w**^⊤^**x** + *b*) model formulations. One would expect the affine models to fit the data better, but the difference in performance is negligible. This is likely due to the relatively high dimensionality of the datasets, since separating hyperplanes are quite easy to find in high dimensions whether or not they are restricted to pass through the origin. We expect a larger performance difference on datasets with smaller feature dimensions (< 10).

**Figure S8:**
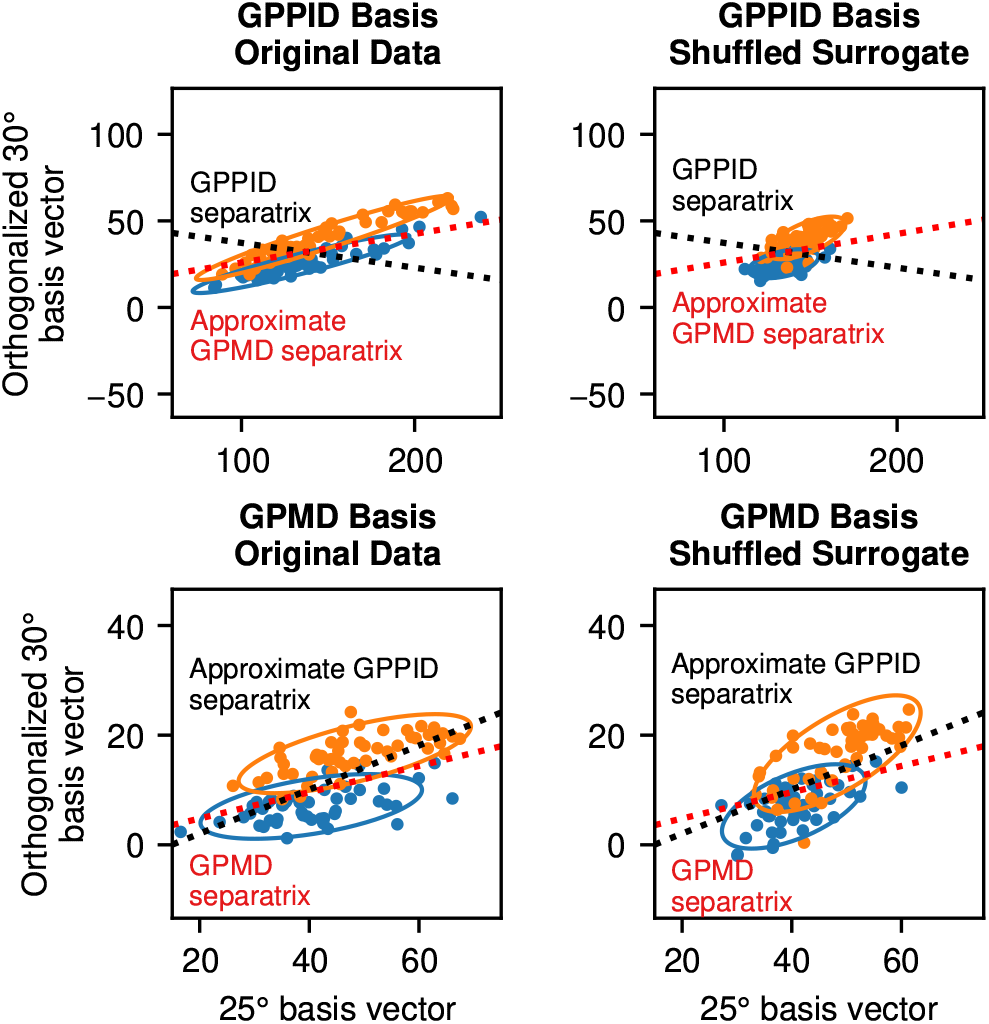
The same visualization as in Figure 7A, but with a surrogate correlation-free dataset created by shuffling each neuron’s responses conditional on the class label. The conclusions remain the same: the data display significant correlations in the GPPID basis, making the GPPID separatrix a poor decision boundary. The GPPID separatrix only performs well when correlations are removed from the dataset (in this case, using shuffling). In the GPMD basis, the difference between the correlated and shuffled surrogate datasets is much less pronounced, indicating that the GPMD’s projection decorrelates the data somewhat.

**Figure S9:**
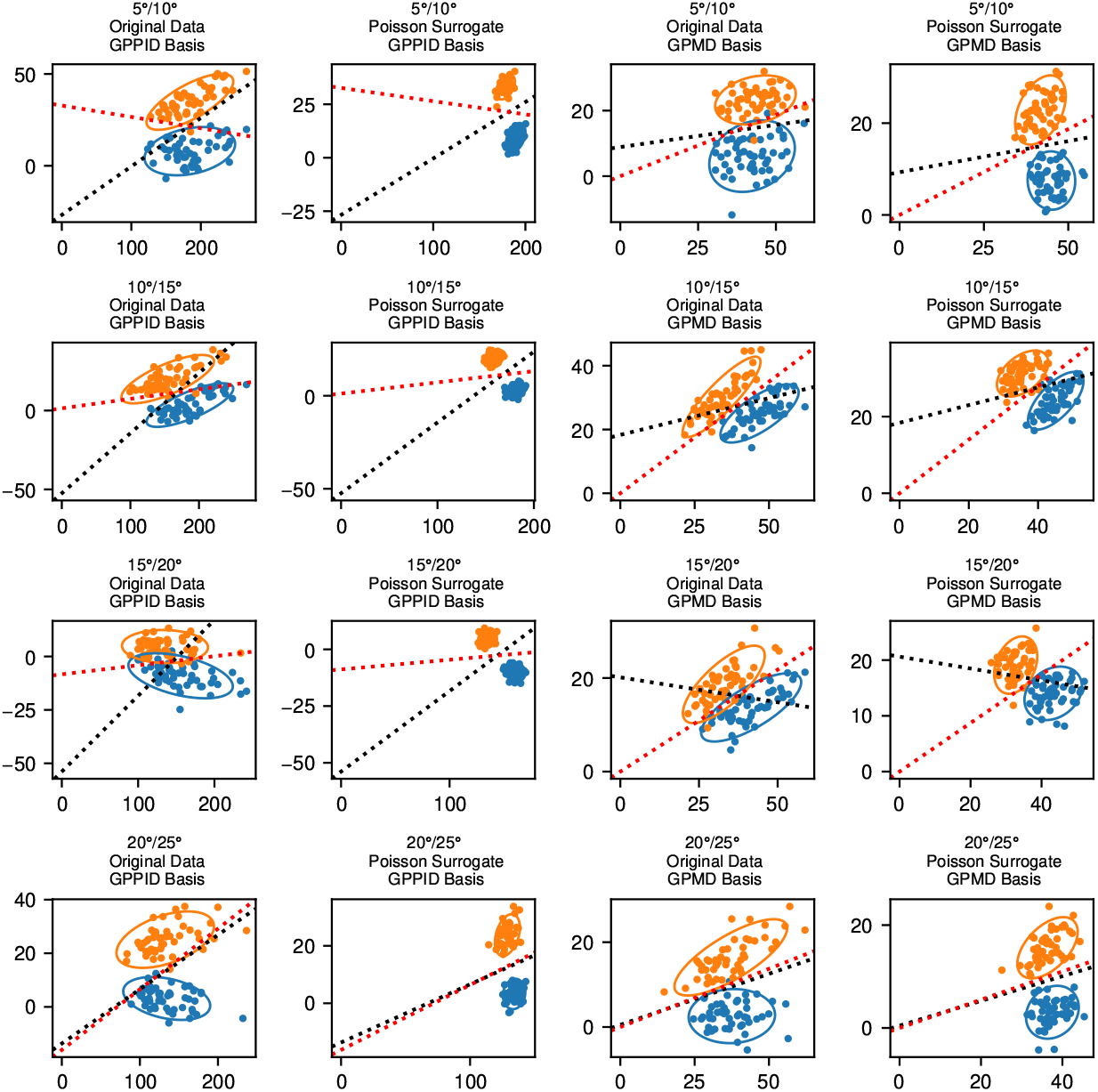
The same visualization as in Figure 7A, for an additional four neighboring stimuli pairs. The GPMD separatrices are colored red, and the GPPID separatrices are colored black. In each case, the general patterns hold: the GPPID fails to separate the correlated data but performs well on the uncorrelated surrogate dataset, the GPMD performs reasonably well in all cases, and the GPMD basis appears to decorrelate the data somewhat.

**Figure S10:**
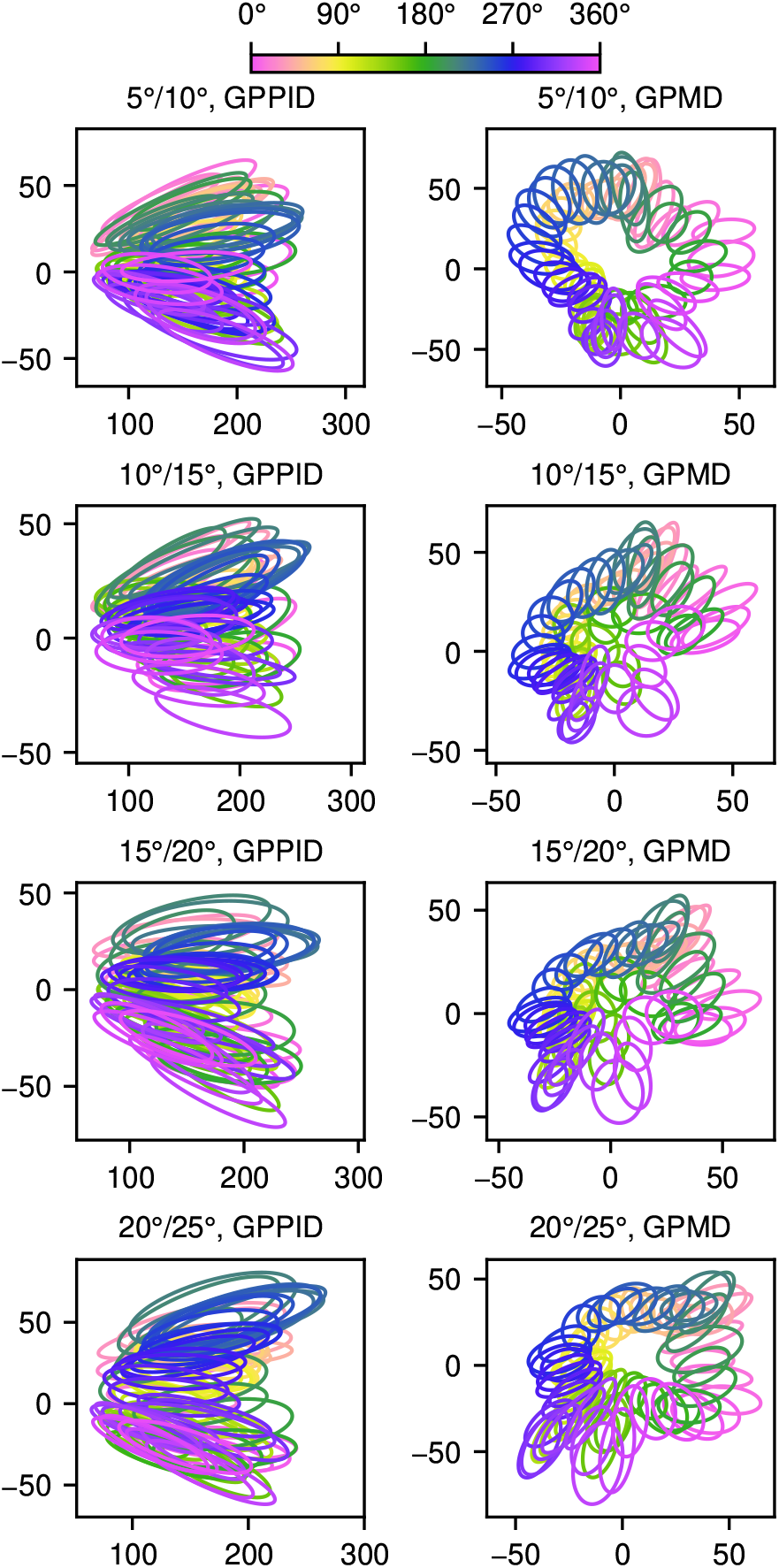
The same visualization as in Figure 7B, for an additional four neighboring stimuli pairs. In each case, the general pattern holds: the GPMD basis produces far superior class separation. In general, the GPMD maps the classes to lie on an ellipse with a 180° period.

**Figure S11:**
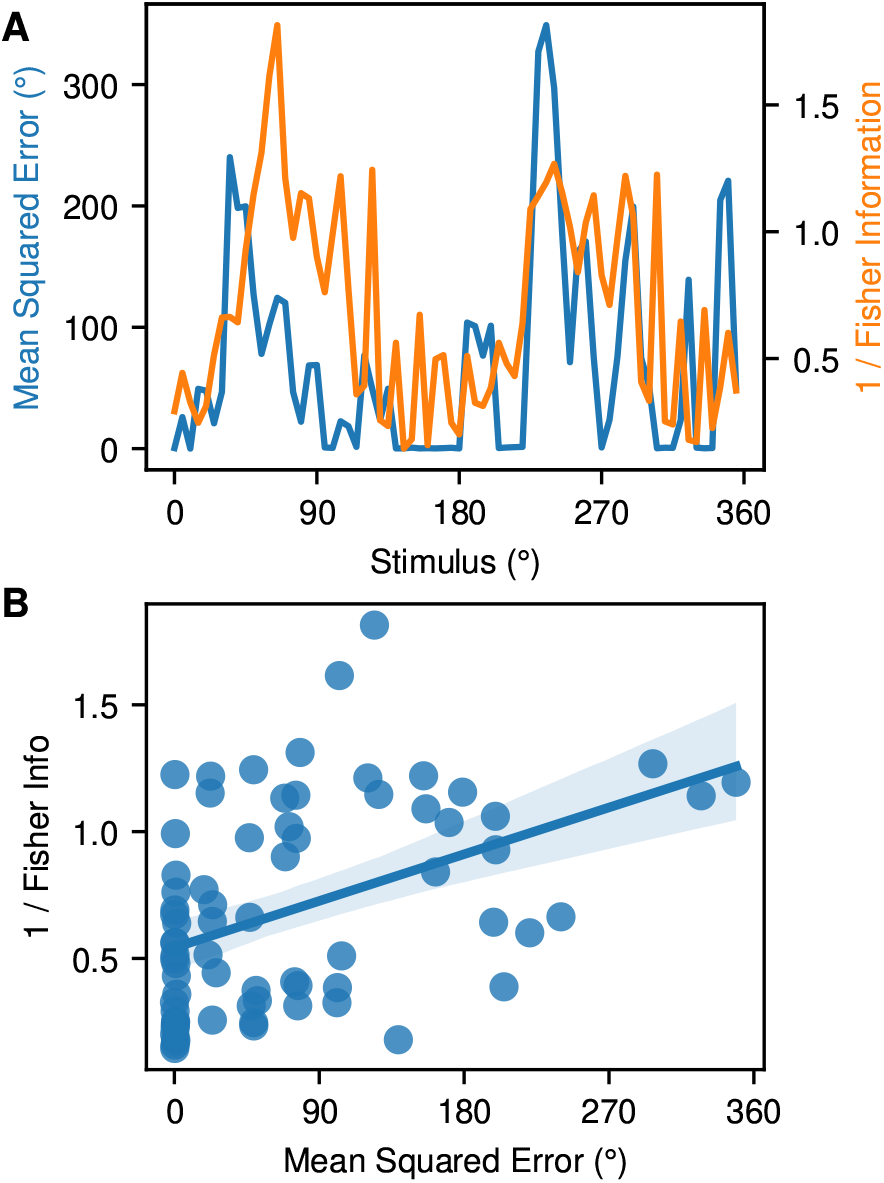
A visualization of how neural selectivity influences decoder performance. We quantified neural selectivity using the total Fisher information, estimated by summing the Fisher information across neurons, using the third monkey dataset. Assuming a Poisson response distribution, the Fisher information for each neuron as a function of the stimulus is the derivative of its tuning curve squared divided by the mean, *f*′(*θ*)*f*(*θ*) (Seung & Sompolinsky, 1993). The inverse of the total Fisher information bounds the minimum achievable mean squared error (Abbott & Dayan, 1999). We wished to compare the inverse total Fisher information of the neural population to the mean squared error (MSE) achieved by the GPMD decoder. Panels **A** and **B** show two views of this information: **A** compares MSE and inverse Fisher information for each stimulus orientation, and **B** is a scatter plot comparison that demonstrates that MSE and inverse Fisher information are positively correlated (linear regression line and 95% confidence interval). The inverse total Fisher information does not lower bound the MSE, since our Fisher information estimate assumes uncorrelated noise. Nevertheless, lower inverse Fisher information is associated with lower MSE, showing that increased neural selectivity contributes to lower decoding error.

